# A benchmarking study of SARS-CoV-2 whole-genome sequencing protocols using COVID-19 patient samples

**DOI:** 10.1101/2020.11.10.375022

**Authors:** Tiantian Liu, Zhong Chen, Wanqiu Chen, Xin Chen, Maryam Hosseini, Zhaowei Yang, Jing Li, Diana Ho, David Turay, Ciprian Gheorghe, Wendell Jones, Charles Wang

## Abstract

The COVID-19 pandemic is a once-in-a-lifetime event, exceeding mortality rates of the flu pandemics from the 1950’s and 1960’s. Whole-genome sequencing (WGS) of SARS-CoV-2 plays a critical role in understanding the disease. Performance variation exists across SARS-CoV-2 viral WGS technologies, but there is currently no benchmarking study comparing different WGS sequencing protocols. We compared seven different SARS-CoV-2 WGS library protocols using RNA from patient nasopharyngeal swab samples under two storage conditions. We constructed multiple WGS libraries encompassing three different viral inputs: 1,000,000, 250,000 and 1,000 copies. Libraries were sequenced using two distinct platforms with varying sequencing depths and read lengths. We found large differences in mappability and genome coverage, and variations in sensitivity, reproducibility and precision of single-nucleotide variant calling across different protocols. We ranked the performance of protocols based on six different metrics. Our results indicated that the most appropriate protocol depended on viral input amount and sequencing depth. Our findings offer guidance in choosing appropriate WGS protocols to characterize SARS-CoV-2 and its evolution.

## Introduction

The severe acute respiratory syndrome coronavirus-2 (SARS-CoV-2), a novel coronavirus that initially emerged in December 2019 in Wuhan, China^1,2^, causing the coronavirus disease of 2019 (COVID-19), has led to a pandemic with >45 million confirmed cases and more than 1,200,000 deaths worldwide as of November 1, 2020 (World Health Organization website https://www.who.int/emergencies/diseases/novel-coronavirus-2019). SARS-CoV-2 can be rapidly transmitted from person-to-person, even during the asymptomatic stage^3^, which is challenging healthcare systems and public health response.

The whole-genome sequencing (WGS) of SARS-CoV-2 has been used as a powerful tool to study COVID-19 since the first sequence was released on Jan 10, 2020^4^. Analysis of the SARS-CoV-2 genome allows for understanding the clinical outcome^5^, developing diagnostics^6^ and vaccines^7^ for COVID-19; and enables the tracking of the evolution^8^ and spread of the virus by phylogenetic analysis^9^, which can reveal the dynamics of subtype evolution. To uncover the complete or near-complete sequence of SARS-CoV-2, leading laboratories have used several sequencing protocols, including shotgun metagenomic approaches^10,11^, target-capture sequencing using Twist custom target enrichment^12^, and target whole-genome amplification sequencing by an multiplex ARTIC primer set^13,14^. However, large variations in performance, e.g., genome coverage and single-nucleotide variant (SNV) detection, occur across different protocols. There is no benchmark study that has compared different protocols using the same patient samples, particularly evaluating factors such as variation in viral input, sequencing platform and depth, sample quality and storage condition. Notably, SARS-CoV-2 WGS requires a viral RNA isolation from the clinical samples for sequencing library construction, and there can be orders-of-magnitude differences in viral load across different subjects. A large proportion of clinical samples contain extremely low viral copy number, which may impact the quality of WGS and the confidence in calls of SNV or indel detection.

We report a benchmarking study on SARS-CoV-2 WGS using clinical nasopharyngeal swab samples. We compared seven different library construction protocols and specifically evaluated the cross-protocol performance in sequencing read mappability, viral genome coverage percentage and uniformity, effect of sequence depth, SNV calling concordance (reproducibility), precision (positive predictive value), and sensitivity (proportion of consensus variants identified at different sequencing depths and viral copy number inputs) across protocols. We constructed libraries using RNA samples isolated from either freshly prepared nasopharyngeal swabs in AVL buffer or frozen nasopharyngeal swabs preserved in AVL buffer to evaluate the impact of storage conditions on the protocol performance. Our findings offer guidance in choosing the most suitable SARS-CoV-2 WGS protocols to better associate SARS-CoV-2 variation with its epidemiological and clinical characteristics.

## Results

### 1. Study design, sample characteristics and SARS-CoV-2 WGS library construction and sequencing, sequencing QC and mapping

All samples were obtained directly from eight COVID-19 patients at Loma Linda University (LLU) Medical Center, and the patients’ clinical characteristics and sample information are presented in **Suppl. Table** The study was approved by the LLU Institutional Review Board (IRB #: 5200127). All SARS-CoV-2 viral samples were collected using nasopharyngeal (NP) swabs. To examine the impact of sample storage condition on the performance of SARS-CoV-2 viral WGS, we used RNA from the NP swab samples stored under two different conditions: RNA isolated either immediately from freshly obtained samples or a frozen (−80°C) NP swab preserved in Qiagen AVL buffer (four subjects in each group). We quantified the SARS-CoV-2 virus copy number for all eight clinical samples using SYBR green qRT-PCR (**Suppl. Table 2**).

We compared seven SARS-CoV-2 WGS protocols, and constructed libraries using low (1,000 SARS-CoV-2 viral copy number, referred as 1K) and high viral copy inputs (250,000 and 1,000,000 SARS-CoV-2 viral copy number, referred as 250K and 1M, respectively) (**Fig. 1**, **Suppl. Table 3)**). Protocol 1 (P1) is the QIAseq SARS-CoV-2 Primer Panel V1 (Qiagen) target whole-genome amplification of SARS-CoV-2 using the multiplex ARTIC v3 primer set. Protocol 2 (P2) is the QIAseq FX Single-cell RNA-seq library kit (Qiagen) coupled with human rRNA depletion. Protocol 3 (P3) is the QIAseq FX Single-cell RNA-seq library kit (Qiagen) coupled with both human and bacterial rRNA depletions. Protocol 4 (P4) is the Tecan Trio RNA-seq kit (NuGEN) coupled with human rRNA depletion and utilized single primer isothermal amplification (SPIA) technology for SARS-Co-V-2 amplification. Protocols 5 and 6 (P5 and P6, respectively) refer to an in-house cDNA synthesis recipe with a mix of random primers, oligo(dT), and four pairs of SARS-CoV-2 specific primers, followed by using either the Illumina DNA library preparation kit-DNA Nano (P5) or the Nextera XT (P6) kit. More recently, Qiagen released its modified SARS-CoV-2 Primer Panel kit (V2) which we also included as Protocol 7 (P7) and benchmarked it with the other six protocols.

**Figure 1.**
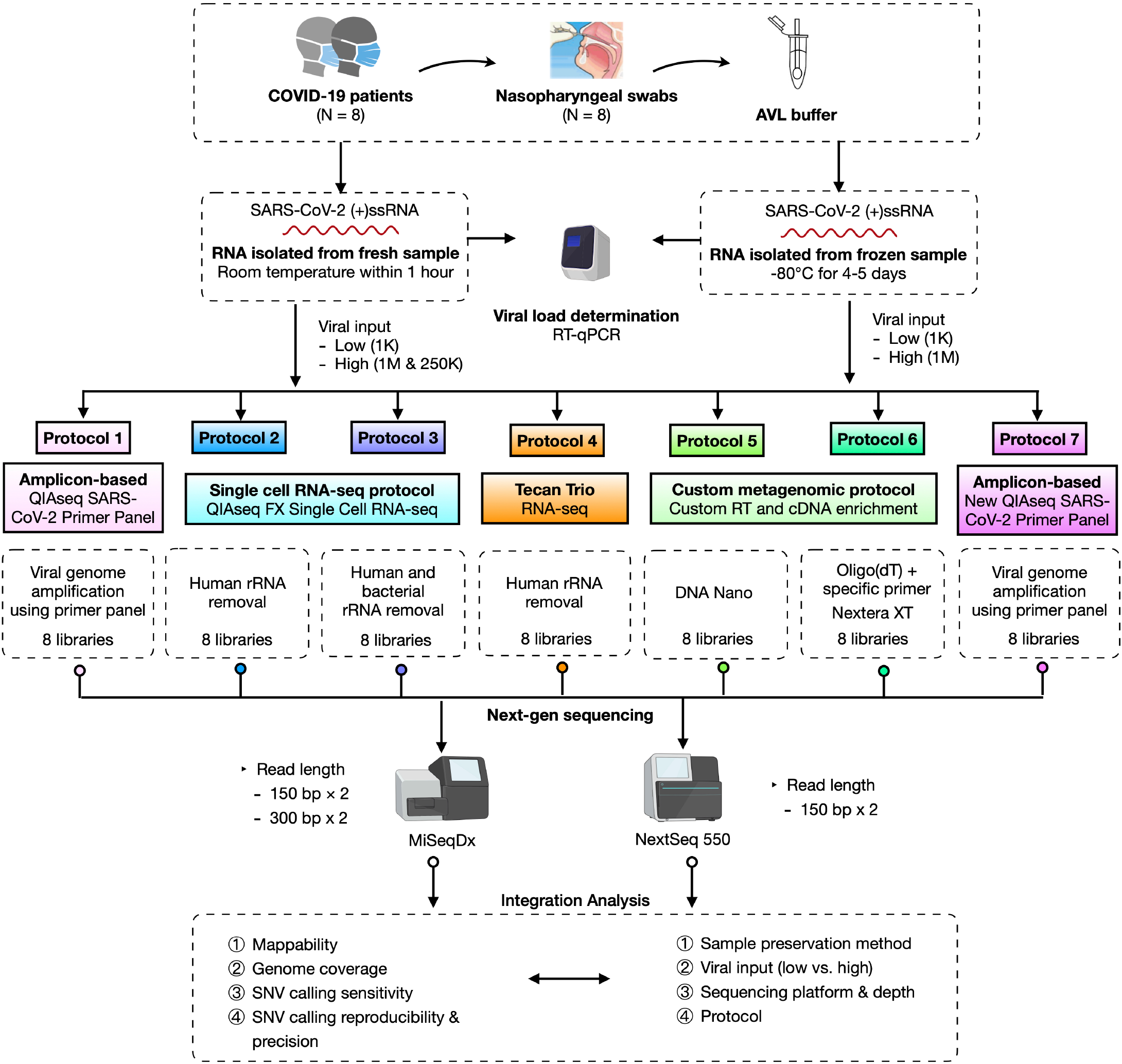
Schematic overview of the experimental design and workflow. Eight COVID-19 positive patient nasopharyngeal swab samples were used to construct the SARS-CoV-2 whole genome sequencing (WGS) libraries using seven protocols. Two different sample storage conditions were compared. For fresh samples, three different viral inputs, i.e., 1000 (1K, low) vs. either 250,000 or 1 million (250K or 1M, high) SARS-Co-V-2 viral copies, were used from each same sample, whereas for frozen samples, two different viral inputs, i.e., 1000 (1K, low) vs. 1 million (1M, high) SARS-Co-V-2 viral copies from each same sample, were used. P4 used different samples at low input vs. high input due to minimal total RNA amount required. The performances of protocols were benchmarked based on viral input, sequencing platform and depth, mappability, viral genome coverage and coverage uniformity, and sensitivity, reproducibility, as well as precision across seven protocols.

Overall, 56 libraries were generated across seven protocols (**Suppl. Tables 3&4**). Of these, 36 libraries were sequenced on both MiSeqDx (300×2 bp and 150×2 bp, paired-end) and NextSeq 550 (150×2 bp, paired-end) (**Suppl. Table 4**). The sequencing reads were mapped to SARS-CoV-2, human, and bacterial reference genomes. Overall, there was no substantive difference in the mapping rates between the MiSeqDx and NextSeq 550 platforms (**Suppl. Fig. 1, Suppl. Tables 5&6**).

### 2. Sequence mapping to SARS-CoV-2 viral, human, and bacterial genomes

We compared the reads mapped to the SARS-CoV-2, human, and bacterial reference genomes between fresh and frozen samples across seven protocols **(Fig. 2a-c)**. We observed statistically significant differences in mapping rates to the viral genome between fresh and frozen samples (p<0.0001 —frozen samples generally had better viral mapping) after accounting for protocol, viral load, and input amount. However, we also observed that the lower mapping rate could be improved and compensated by deeper sequencing (i.e., ~2x deeper for fresh samples vs. frozen) for certain protocols (**Fig. 2a**). The significant differences in viral mapping were more easily seen in P1, P2, P3, and P4 generally corresponded to higher off-target bacterial sequence observed in fresh versus frozen samples (**Fig. 2a, c**). One exception was the P1 for which fresh samples also resulted in higher (off-target) human mapping rates (**Fig. 2b**). Overall, the above results suggested that frozen samples performed better than or equivalent to fresh samples in their mappability and on-target percentage to the SARS-CoV-2 genome for all protocols with P7 having very high mappability with either sample preservation storage methods (**Fig. 2a-c**).

**Figure 2.**
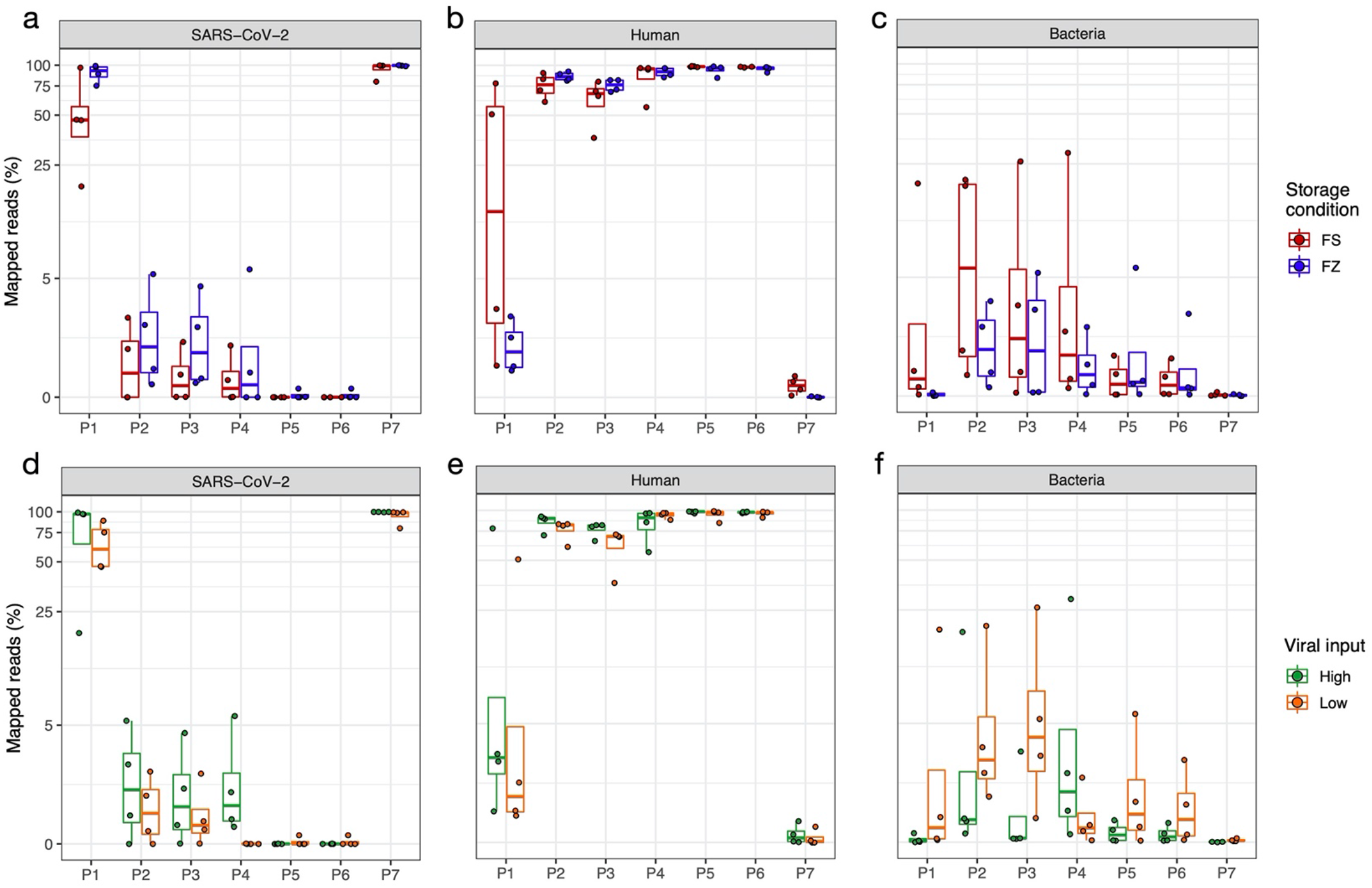
Comparison of reads mapped to SARS-CoV-2, human, bacterial genomes. The top panel box plots show the percentages of reads mapped to (**a**) SARS-CoV-2 genome, (**b**) human genome, and (**c**) bacterial genome, at two different sample storage conditions, i.e., fresh (FS) vs. frozen (FZ). The bottom boxplots show the percentages of reads mapped to (**d**) SARS-CoV-2 genome, (**e**) human genome, and (**f**) bacterial genome at different viral input, i.e., high (250K or 1M) vs. low (1K). Y-axis shows the percentages of reads (log10 scaled, only for panels **a** and **d**); X-axis shows the protocol number. FS (fresh): RNA isolated from fresh samples; FZ (frozen): RNA isolated from frozen samples; Low: 1K viral input; High: 1M and 250K viral input.

Read mappability to the SARS-CoV-2 viral genome versus human and bacterial genomes were clearly different across protocols regardless of RNA prepared from fresh or frozen samples. The ARTIC amplicon-based target genome amplification technology P7 had the highest read mapping percentage (96.9% ± 7%) to the SARS-CoV-2 viral genome with the lowest reads mapping percentage to human genome (0.17% ± 0.2%) among the seven protocols (**Fig. 2a**). However, P1, an earlier version of the ARTIC amplicon-based protocol, had the second highest read mapping percentage (71% ± 30.2%) to the SARS-CoV-2 viral genome with a substantially higher mapping rate (17.4 ± 29.9%) to the human genome as compared to P7 (**Fig. 2a**). All the other metagenomic approach-based protocols had starkly fewer reads mapped to the reference viral genome (< 5.7%). For the QIAseq FX Single-cell RNA-seq library kit incorporated with human ribosomal RNA depletion (P2), the viral mapping rates were 1.6% ± 1.8%, and the mapping proportions to human genome were 80.9% ± 10.7%. When the same library kit was incorporated with both human and bacterial ribosomal RNA depletions (P3), the viral mapping rates dropped slightly to 1.2% ± 1.5%, and the mapping rates on the human genome also dropped to 69.4% ± 14.6%. Both P5 and P6 had the lowest percentage of reads mapped to the SARS-CoV-2 viral genome (0.03% ± 0.08%), and the highest mapping rates to human genome (96.1% ± 5% and 96.4% ± 2.5%, respectively) (**Fig. 2a**). We also noticed that P2 had the highest rate of reads mapped to bacterial genome overall (5.9% ± 8.3%) among the seven protocols (**Fig. 2c**).

To evaluate the variability of viral load on the SARS-CoV-2 WGS performance, we compared the SARS-CoV-2, human, and bacterial reference mapping rates between low (1K copies) and high viral inputs (250K and 1M copies) across seven protocols (**Fig. 2d-f**). In our study design, different levels of SARS-CoV-2 viral inputs were generated using either original undiluted patient NP sample-derived RNA or a dilution from the identical higher viral load samples across all protocols (**Suppl. Table 3**). We found that high viral inputs (250K and 1M copies) had a higher mapping rate to the SARS-CoV-2 genome compared to low viral inputs across all protocols as expected, although the difference is smallest for P5, P6 and P7 [high vs. low, diff= 0.38, p = 0.01, generalized linear mixed model (glmm, statistical testing table now shown), **Fig. 2d**)]. P1 and P7 performed well with P7 having better mapping rates than P1 at either input level (high viral input vs. low viral input mapping rates: P7, 99.7% ± 0.3% vs. 94.2 ± 9.7%) with significantly less sequencing reads mapped to either human or bacterial genomes (**Fig. 2d-f**), while P5 and P6 performed very poorly for both low and high viral inputs with extremely low SARS-CoV-2 viral mapping rates compared to other protocols (**Fig. 2d**). For P4, low viral input resulted in orders of magnitude lower SARS-CoV-2 viral mapping rates compared to high viral inputs, e.g., 0.003% ± 0.01% vs. 2.1 ± 2.5%, suggesting P4 would require a higher viral input to obtain adequate SARS-CoV-2 viral genome coverage when sequencing depth is limited (**Fig. 2d**). In general, the mapping rates to SARS-CoV-2 differed significantly by protocol (p< 2.20E-16, glmm), and by sample storage condition (p=2.45E-09, glmm). According to the differences in their least square means, P7 had a statistically significantly higher SAS-CoV-2 mapping rate compared to the other six protocols (P7 vs. P1, diff = 1.27, p = 1.52E-05, glmm) followed by P1 (P1 vs. P2, diff = 3.25, p <2.20E-16, glmm, statistical testing table now shown). The advantage in mapping rate of P7 and P1 is directly attributable to their viral amplicon-based design. Protocols P2-P4 were less easily differentiated by mapping rate (**Fig. 2d**).

### 3. Viral genome coverage and the effect of sequence read depth

To determine the impact of read depth to the SARS-CoV-2 genome coverage, we down-sampled all of the library-run datasets to 50,000 (50K), 100,000 (100K), 500,000 (500K), 1,000,000 (1M), 5,000,000 (5M), 10,000,000 (10M), 15,000,000 (15M), and 20,000,000 (20M) paired-end (PE) reads, and evaluated the SARS-CoV-2 viral genome coverage at different sequence depths across seven protocols (**Fig. 3**). We noticed one sample (NP12) performed extremely differently compared to other samples across protocols (green line in **Suppl. Fig. 2**). Therefore, we excluded this sample from the analysis for **Fig. 3**. Both amplicon-based protocols P1 and P7 achieved significantly higher coverage for the SARS-CoV-2 at a threshold of >10X reads at each base (termed min10X), even with comparatively lower overall read depths compared to other protocols. Particularly, P7 consistently had higher percentages of SARS-CoV-2 genome coverage (min10X) regardless of viral input amount (**Fig. 3a-b**). At low viral input but with only 1M sequencing reads, P7 had 94% ± 5.1% of the SARS-CoV-2 viral genome coverage (min10X). By comparison, P1, the other amplicon-based target genome amplification protocol, could only achieve 71% ± 23.7% of the viral genome coverage (min10X) at a sequencing depth even with 5M reads (**Fig. 3a**). Interestingly, at low viral input, P2 and P3 achieved higher percentages of genome coverage (min10X) than P1 at 5M reads (**Fig. 3a**). We found that P2 and P3 achieved almost complete coverage of SARS-CoV-2 genome (min10X, 95.6 ± 2.3% and 93.8 ± 5.2%, respectively) with 5M reads at low viral input (**Fig. 3a**). P4 - P6 performed poorly with low viral input even at 15M read depth. However, the performance of viral genome coverage (min10X) at the low input generally was not always consistent with what was observed at the high input for each protocol except for P2 which achieved nearly 100% genome coverage (min10X) at the low input with 10M reads, and it reached to nearly 100% genome coverage (min10X) at high input with 1M reads (**Fig. 3a-b**).

**Figure 3.**
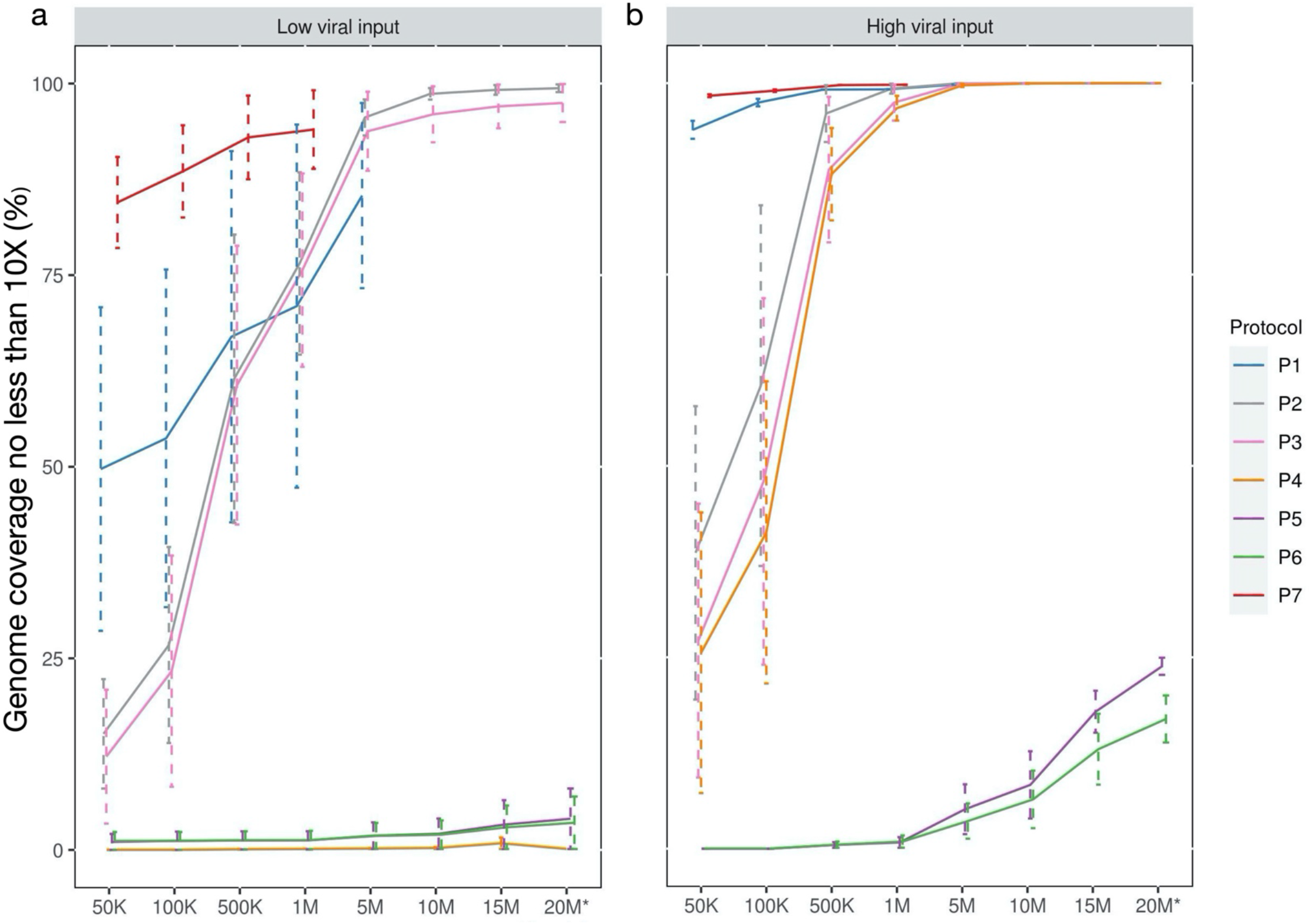
SARS-CoV-2 genome coverage across seven protocols at different sequence depths. The average genome coverage of samples at (**a**) low viral input (1K) and (**b**) high viral input (250K or 1M) with different down-sampled read depths. Bars represent the standard errors. The breath of coverage of SARS-CoV-2 genome was defined as the percentage of SARS-CoV-2 reference genome for which the genomic locations (bases) had minimal10X coverage. Seven down-sampled read depths (from 50K to 15M) plus the original total reads (20M*) for each sample were provided. *For samples with sequencing reads more than 20M reads, the data only showed 20M down-sampled reads. The sequencing down-sampling were performed using seqtk (v1.0.r75) with ‘sample’ command. Sample NP12 was excluded in this figure.

Furthermore, when the samples contained high viral input, both P1 and P7 could achieve high viral genome coverage (min10X) at lower read depths as compared to all other protocols (**Fig. 3b**). For example, when the sequencing depth was at only 50K PE, P1 had 93.9% ± 1.2% of genome coverage whereas P7 reached 98.3% ± 0.2% of the SARS-CoV-2 genome coverage (min10X, **Fig. 3b**). In contrast, at high viral input, three RNA-seq based metagenomics protocols P2, P3 and P4 had similar genome coverage levels across different sequencing depths (e.g., min10x at 1M read depth: P2, 99.3 ± 0.6%; P3, 97.5 ± 2.4%; and P4, 96.7 ± 1.6%, respectively, **Fig. 3b**). Nevertheless, for P5 and P6, the observed genome coverage (whether >1X or >10X) was consistently low at even >20M reads, regardless of high or low viral input (**Fig. 3a-b**), suggesting that P5 and P6 would not be suitable in practice to achieve necessary coverage for SNV detection. Notably, when all the samples (including NP12) were included for genome coverage analysis, we observed a similar relative pattern of performance by protocol between viral input levels as demonstrated in **Fig. 3** but with greater variation (**Suppl. Fig. 3a-b**).

### 4. SARS-CoV-2 viral genome coverage uniformity comparisons

To evaluate the coverage quality and regional bias relative to the viral reference genome between protocols, we computed the average coverage by genomic position across all samples across all protocols at a depth of 5M total reads (**Fig. 4a-b**). As expected, due to their amplicon-based nature, P1 and P7 had the highest average coverage compared to all the other protocols (**Fig. 4a-b**). A great majority of the regions across the SARS-CoV-2 genome had ~1,000X coverage while some regions were near or exceeded 10,000X coverage for P1 and P7 at high viral input (**Suppl. Fig. 4**). The variations or “spikes” of the coverage in many regions across the viral genome were much more pronounced at the low viral input as compared to the high viral input derived from the same samples (**Suppl. Fig. 5**). For P2 and P3, the average coverage across the entire viral genome usually ranged from 300-600X (**Fig. 4a**), whereas for P5 and P6, only certain regions were sequenced, and many regions showed no coverage in the samples with either low or high viral input (**Fig. 4a-b**). Furthermore, we observed that protocols P2 and P3 had much higher coverage at the 3’ end of the viral genome, which was likely introduced by oligo(dT) primers during cDNA synthesis. Other than the 3’ end of the viral genome, P2 and P3 had comparably even coverage across the viral genome regardless of viral inputs (**Fig. 4a-b**). However, the viral genome coverage was significantly different between the low and high viral input in P4 which had scarcely or no coverage across whole genome at low viral input (**Fig. 4a-b**). At high viral input, P4 had excellent coverage uniformity across the whole genome (**Fig. 4b**). Finally, for completeness, in addition to greatly reduced viral genome coverage for P5 and P6, but there was also a lack of coverage uniformity regardless viral input (**Fig. 4a-b**).

**Figure 4.**
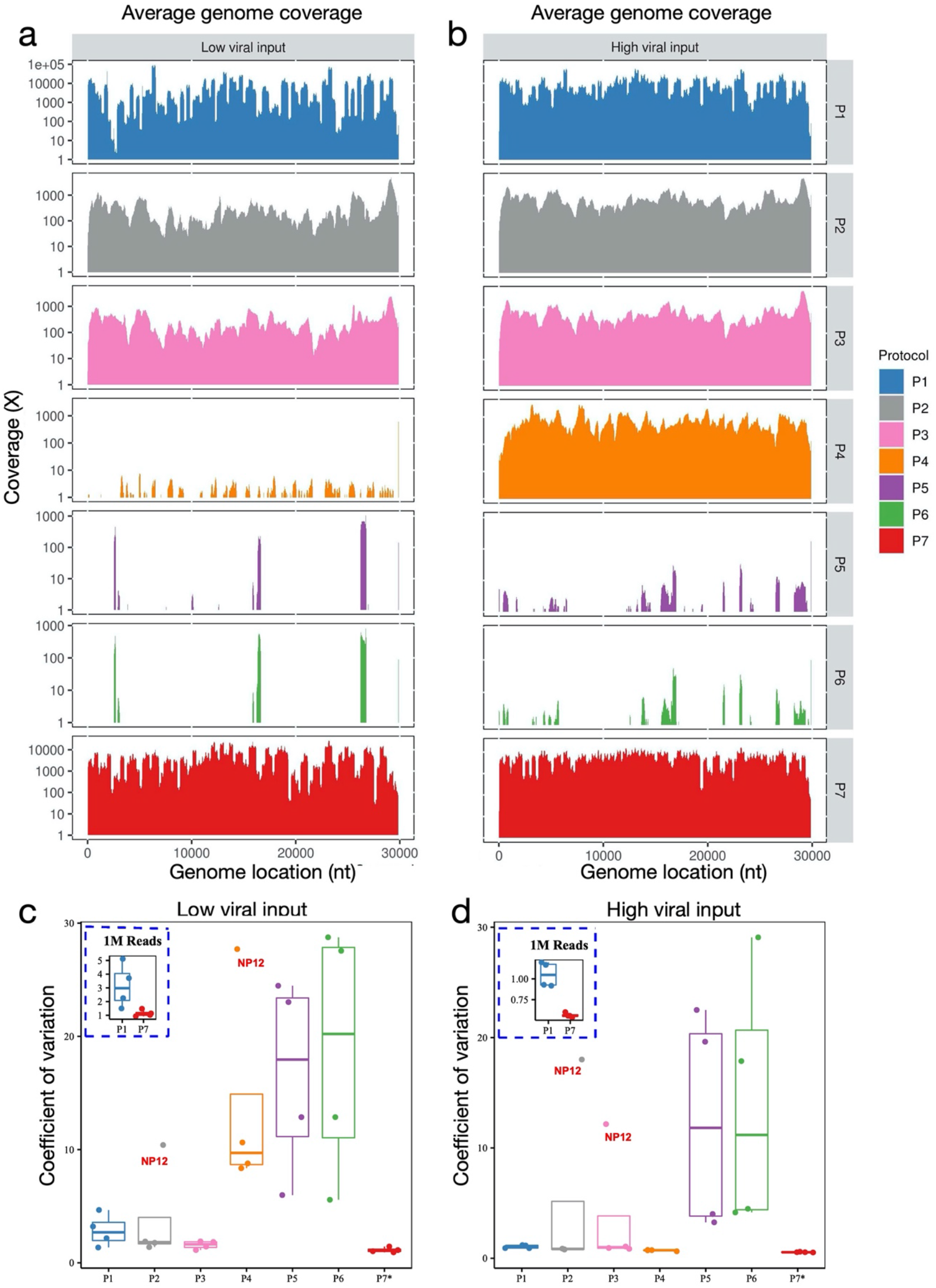
SARS-CoV-2 Genome coverage uniformity across seven protocols. **(a)** Average of SARS-CoV-2 genome coverage based on four samples at low viral input (1K); (**b**) Average of SARS-CoV-2 genome coverage based on four samples at high viral input (250K or 1M); for P1-P6, all the sequencing data were down-sampled to 5M reads. For P7, all the sequencing data were down-sampled to 1M reads. The boxplots in the panels (**c)** and (**d)** show the mean (M) with 2X standard deviation (SD) of coefficient of variation (CV) of the SARS-CoV-2 genome coverage uniformity at low viral input (**c**), or at high viral input (**d**). CV metric was computed using the SD and M of the coverage at each reference genome position. The inserted dash line square shows the boxplots of CV (M ± SD) for P1 vs. P7 at 1M read depth for low viral input (**c**) or high viral input (**d**). *P7 only had sequencing data available at 1M read depth, thus only 1M reads of sequencing data was used to calculate the CV.

We further compared the coverage uniformity using a quantitative metric, i.e., coefficient of variation (CV) across seven protocols (**Fig. 4c-d**) computed using the averaged coverage depth by reference genome position. Amplicon-based P7 had the best uniformity of genome coverage at both low and high viral inputs and was the only protocol with a coverage CV at or less than 1 (i.e., 100%, **Fig. 4c-d**), whereas protocols P1-P3 had ~ 3-4 times higher CV compared to P7 at either viral inputs (**Fig. 4c-d**). However, for P4 at high viral input, consistent with what was demonstrated in the coverage tracks (**Fig. 4a-b**), the CV was very small (~1, most similar to the P7, **Fig. 4d**), indicating excellent uniformity in genome coverage. Conversely, at low viral input, P4 had much larger CV values (range of [9,18], **Fig. 4c**). We also examined the relative impact of viral input (high vs. low, pair-wise, derived from the same clinical sample) on coverage uniformity using our CV metric (**Suppl. Fig. 6**). We found that the uniformity and overall coverage of the SARS-CoV-2 genome as evaluated by CV improved remarkably for each of six protocols (P4 was not evaluated due to distinct clinical samples used for low vs. high input), when higher viral inputs (e.g., 250K or 1M vs. 1K copies) were used (**Fig. 6a-b**). Particularly, the coverage uniformity for P3 was the least affected by viral input changes. Coverage uniformity of P2 and P7 were also less impacted by low versus high viral input levels than other protocols (**Suppl. Fig. 6b**).

To gain a deeper understanding on the variations in read depth in certain regions and the differences of these variations between the two related amplicon-based protocols (i.e., P1 and P7), we compared the genome coverage profiles of three samples with high viral input (1M copies) for which the WGS libraries were constructed using P1 and sequenced at 5M read depth (**Suppl. Fig. 7**). We noticed that all three samples shared similar coverage patterns across the whole SARS-CoV-2 genome, suggesting that those local high spikes were primer-set dependent. We examined the primer sets corresponding to those highly variable regions, and found that four mostly over-represented spiking regions in the three samples using P1 were associated with about 20 primer sets whose amplified genome regions were covered by multiple amplicons (**Suppl. Fig. 7**). Although the ARTIC V3 primer set was designed to cover each genome position with two amplicons (with the exception of the regions covered by amplicons 1 and 98), high coverage regions were associated with three or more amplicons. Consistent with this fact, the coverages for those regions were roughly equal to the sum of the coverages from three to four individual amplicons, supposing each amplicon had a similar amplification efficiency. Furthermore, at least one extra alternative primer pair was linked to each high coverage region, hinting a redundancy for some ARTIC V3 primers. In contrast, our study showed that P7 had a more uniform coverage across all regions compared to P1 (**Fig. 4a-d**, **Suppl. Figs. 4 & 5**), even though there was no change made to the ARTIC V3 primers except for some modification of other related reagents in the P7 kit by Qiagen (proprietary, personal communication with Qiagen). In addition, for P1, our data suggested that the relatively low coverage in other regions, i.e., 1770-3303 and 23609-24856, might be due to the low efficiency of primers (**Suppl. Fig. 7**). Finally, it was unsurprising to observe low coverage at both the 5’- and 3’-regions for P7 and P1 since only one primer pair covered each of these regions.

### 5. Sensitivity of viral genome variant identification using consensus SNVs, viral subtypes and phylogenetic analysis

Genome variants in clinical samples were called from BAM files after removing duplications and primer sequences from amplicon reads. In order to evaluate the accuracy of variant calling, we called variants from clinical samples prepared from P1, P2, P3, P4, and P7 using VarScan 2 (v2.4.4)^15^ and iVar (v1.2.2)^16^ against the reference SARS-CoV-2 genome NC_045512.2. The putative SNVs were defined as variants with 10X minimum coverage and > 80% allele frequency, the default setting for VarScan. Our WGS data showed that five out of the eight clinical samples were able to produce SNV calls with the other three samples failing due to insufficient coverage. We analyzed the results from the independent preparation and library methods and found certain SARS-CoV-2 viral SNVs recurred across protocols and across patients. In particular, we identified ten SNVs that were found by at least three distinct protocols in at least one patient, all at high allele frequency. Seven of these ten were identified in more than one patient. We termed these ten SNVs as the consensus SNVs for this study, and we established that a consensus SNV for a patient was a consensus SNV from the overall study that was identified by at least two distinct protocols. These ten consensus SNVs, for which our measurements of sensitivity were derived, are provided in **Suppl. Table 7**. Notably, all consensus SNVs in patient samples had high (> 90%) average allele frequency, which was to be expected for a haploid-type genome. Also, of importance, all 16 SNVs in the study that had observed allele frequency above 80% but were not identified as consensus SNVs were: **a**) never replicated across protocols for the same sample; **b**) were never identified in more than one sample; and **c**) were all observed only when using low viral copy input (data not shown). Therefore, higher viral copy input was strongly associated with better cross-protocol SNV reproducibility as discussed in more detail in the next section.

Sensitivity results at high viral input for consensus SNVs are shown for three representative clinical samples (NP08, NP29, NP30) in **Fig. 5a**, and **5b**. All putative SNV data from high viral input at different read depths are provided in **Fig. 5c** while **Fig 5d** illustrates the spatial organization of the consensus SNVs by sample. At high (1M) viral copy input level, 5M 150×2 paired-end reads were sufficient to detect nearly all consensus SNVs in the three representative clinical samples using any of P1–P4 except P7 for which only 1M reads were needed (**Fig. 5b**). Although all five protocols were able to detect the majority of consensus SNVs from as low as 0.5M reads, P1 and P7 exhibited better performance than P2, P3, and P4 at low read depth due to a high percentage of on-target reads. We found that only 0.5M reads were sufficient for P7 to properly detect roughly 95% of the consensus SNVs (**Fig. 5b**). Surprisingly, P1 required 5M reads to detect more than 90% of the consensus SNVs, which was more than expected and contrasted greatly with P7 (**Fig. 5b**). At 5M reads, P2 identified all consensus SNVs in all samples while P3 and P4 had sensitivity levels that exceeded P1 (**Fig. 5a-b**).

**Figure 5.**
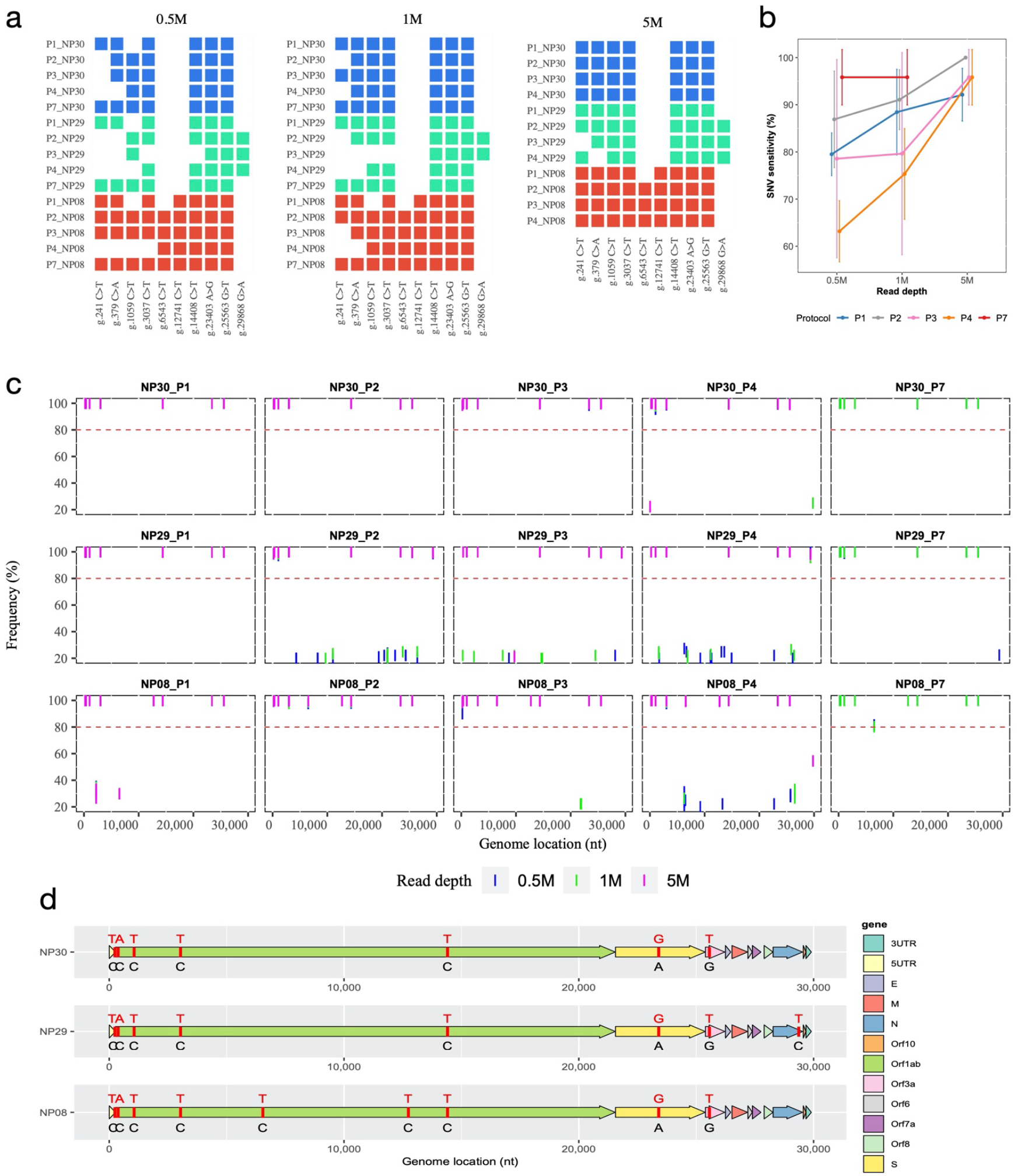
Influence of library protocol and read depth on SNV detection. (**a**) Comparison of SNV calling across P1, P2, P3, P4 and P7 protocols at various sequencing depth; all SNVs with min10X and frequency higher or equal to 80% were deemed as true SNVs and show in rectangle plot. (**b**) Sensitivity of SNV detection at different sequencing depth; only the data at high viral input (250K and 1M) were used and showed. Sensitivity was calculated using the consensus SNVs as the set of positives. Each data point was the average of calling percentage by each protocol (n=3). The calling percentage was calculated as the number of consensus SNVs called by each protocol divided by the total number of consensus SNVs in each sample. (**c**) SNVs called by P1, P2, P3, P4, and P7 at three different read depths (0.5M, 1M, 5M) in three samples at high viral inputs. The red dash line shows the threshold of SNV calling, i.e., frequency higher or equal to 80%. (**d**) Consensus true SNVs depicted based on the NC_045512.2 genome; four sub-genome types were identified from all clinical samples in this study. Reference nucleotides are colored in black and the SNVs are colored in red.

**Figure 6.**
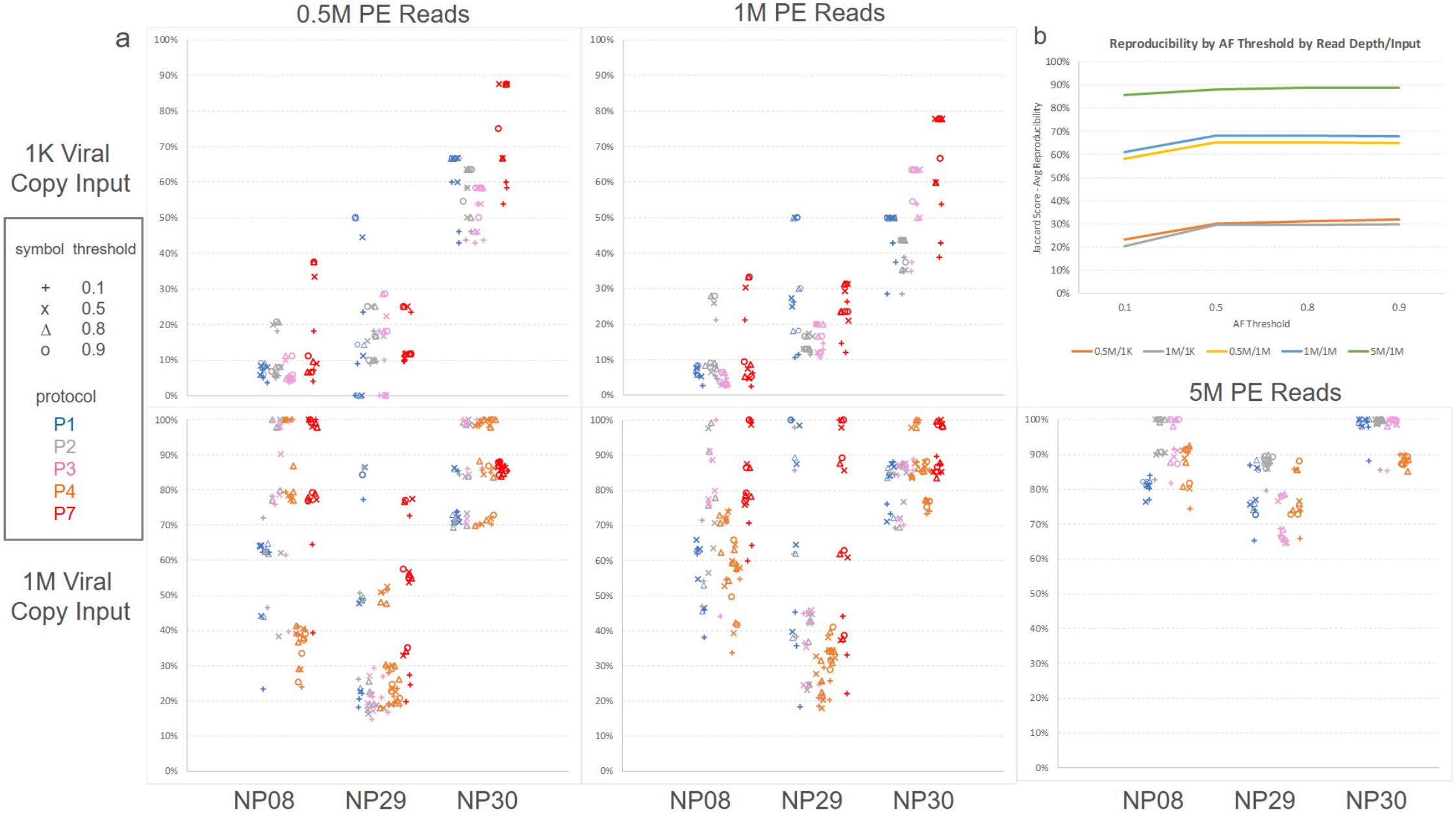
Influence of protocol, sample, read depth, input amount, and allele frequency calling threshold on SNV reproducibility. (**a**) By input amount (1K at the top vs. 1M at bottom) and read depth (0.5M, 1M, 5M), each graph indicates the reproducibility (%) of SNV calls between protocols (P1, P2, P3, P4, P7) for a given sample, where each point represents another protocol with which the indicated protocol (by color) was compared. Only at 1M viral input were the data available for 5M PE reads and showed (bottom panel, right), no data available at 5M reads at low input. X-axis shows the samples (NP08, NP29, NP30), Y-axis shows the SNV calling reproducibility. (**b**) Jaccard score showing the average reproducibility between protocols across all combinations of protocols when varying read depth (0.5M, 1M, and 5M), viral input amount (1K, 1M), and allele frequency calling threshold; X-axis shows allele frequency (AF) threshold, Y-axis shows the Jaccard score—average reproducibility (%); 0.5M/1K: 0.5M reads/1000 viral copies, 1M/1K: 1M reads/1000 viral copies; 0.5M/1M: 0.5M reads/1 million viral copies; 1M/1M: 1M reads/1 million viral copies; 5M/1M: 5M reads/1 million viral copies.

However, at lower viral copy input, we observed that several consensus SNVs were not detected. i.e., either observed at low allele frequency or completely undetected across protocols, which indicated a reduced sensitivity in SNV detection with low viral input (**Suppl. Fig. 8&9**). Particularly, at low viral copy input, we noticed that P4 did not make any SNV call at all due to its inadequate genome coverage. However, P2 achieved excellent sensitivity (almost 100%) at 5M PE reads while P7 was able to achieve ~90% sensitivity on average at 0.5M PE reads even with low viral copy input (**Suppl. Fig. 8&9**).

Phylogenetic network analysis of complete SARS-Cov-2 genomes has been conducted to track the transmission of COVID-19^9^ (. Based on the SNVs identified in our tested samples, we performed phylogenetic analyses to explore the relatedness of the genotypes in our samples with the viral strains spread in the country and the world. We identified four viral genome subtypes containing six to nine SNVs within each genome subtype (**Suppl. Fig. 10**). All four viral subtypes appeared to share six of the ten consensus SNVs, indicating that these genotypes were phylogenetically closely related (**Suppl. Figs. 11a-b & 12**). Using a global phylogeny generated via Nextstrain^17^ (**Suppl. Fig. 11**), the cases from the Loma Linda (CA, USA) were most related to the cohort from MI (USA), with all the five virus samples pertaining to clade 20C (**Suppl. Fig. 12**).

### 6. Reproducibility, precision of SNV detection, and the effect of viral input and sequencing depth

To investigate the influence of viral input, sequencing read depth, and protocol on key quality parameters such as reproducibility and precision, we examined protocols P1, P2, P3, P4, and P7 using samples NP08, NP29, and NP30 which had WGS libraries constructed with both 1K and 1M viral inputs. We defined the reproducibility relative to an allele frequency threshold: i.e., a variant was reproducible between protocols A and B (or between input amounts for the same protocol) if the variant from protocol A’s library had an allele frequency equal to or greater than a threshold for which the variant was also identified by protocol B’s library at any allele frequency. We used the Jaccard index for reproducibility scoring. We observed that the protocol, read depth, viral input amount, and the sample itself impacted the reproducibility between protocols on called SNVs (**Fig. 6a**). The sample itself and the viral input amount had the biggest impact on reproducibility, suggesting that there were characteristics about the sample apart from viral input level that influenced the SNV detections. The allele frequency threshold impact was muted once the threshold exceeds 50%. The average reproducibility across protocols using the Jaccard index was almost 90% when using 1M viral copies for input with reasonable allele frequency thresholds and when sequencing the sample to 5M PE reads (**Fig. 6b**). Lowering either read depth to 1M or 0.5M PE reads or reducing the viral input to 1K copies noticeably reduced reproducibility by at least 20% and sometimes by 50% or more (**Fig. 6b**).

Regarding the precision of SNV calling, we noticed that at 1K viral copy input a large number of low allele frequency SNVs (i.e., 5-80%, false) were putatively identified in each sample by P1, P2, P3, and P7 (P4 not evaluated at low viral copy input due to generally poor genome sequence coverage results), especially in sample NP08 (**Suppl. Fig. 13)**. An increase in read depth did not improve the precision of SNV calling, nor did it reduce the number of low allele frequency SNVs (**Suppl. Fig. 13**). As SARS-Co-V2 is a haploid virus, we assumed that the vast majority of these low frequency putative variants were false SNVs. Consistent with this, none of these putative lower allele frequency SNVs was reproduced across at least three protocols (**Suppl. Fig. 13**). On the other hand, in certain clinical samples (NP08 and NP30), some putative false SNVs with high allele frequency (>80%) were called primarily by amplicon-based protocols (P1 or P7) at low viral copy input (**Suppl. Fig. 8a-c,** SNVs without an asterisk), nevertheless these SNVs were not reproduced even in the same samples with high viral copy input (**Fig. 5a&c**).

In summary, we observed that nonreproducible candidate SNVs tended to have one or more of the following characteristics: 1) they tended to have allele frequency below 80% (94% of observed nonreproducible SNVs); 2) they tended to occur when using low viral input (85% of observed nonreproducible SNVs); and 3) they were observed at local pile-up depths of less than or equal to 50 bases (66% of observed nonreproducible SNVs). Our benchmarking study suggested that low viral copy input severely affected reproducibility of SNV calls as well as sensitivity and precision of consensus SNVs.

## Discussion

The COVID-19 pandemic is causing a global health crisis. By November 2020, over 1.2 million deaths were attributable to COVID-19, and the number is continuously growing (World Health Organization website https://www.who.int/emergencies/diseases/novel-coronavirus-2019). There is an urgent need to better understand and track SARS-CoV-2 to improve the viral detection, tracing of the viral transmission, and the development of effective therapeutic approaches. Generating full-length SARS-CoV-2 sequence through next-generation sequencing (NGS) will allow better understanding of its evolution and enhance the treatment strategies for COVID-19^8,18–21^. Here, we compared seven WGS protocols for SARS-CoV-2 using clinical samples from infected patients, benchmarking the performances of these protocols in several aspects including the sequencing read mappability, genome coverage (percentage and uniformity, minimum sequences required); sample storage condition; effects of viral input, sequencing depth, length and platform; sensitivity, reproducibility and precision of SNV calling and related assay factors (e.g., amount of viral input, sequencing depth and bioinformatics pipeline).

The SARS-CoV-2 is a positive-sense single-stranded RNA virus, which has low stability once RNA enzymes are released after cellular destruction. The quality of virus RNA is critical for the detection and the overall genome sequencing. It has been reported that only 47-59% of the positive cases are identified by RT-PCR, possibly due to loss or degradation of virus RNA during the sampling process^22,23^. Starting the RNA isolation immediately following NP swab sample collection may be ideal to minimize RNA degradation; however, immediate isolation is often impractical especially when involving large cohorts of sampling at different time points. Therefore, we compared the samples isolated from two storage conditions, i.e., RNA isolated either immediately from the freshly prepared NP swabs or from the NP swabs in AVL buffer that were frozen at −80C for 5-6 days. We found that although there were differences in the genome mappability between fresh and frozen samples across the protocols where the frozen samples performed slightly better than or equivalent to fresh samples in their on-target percentage to the SARS-CoV-2 genome (**Fig. 2a-c**), there was no practical difference in the on-target sequence mappability for P7. Furthermore, for other well-performing protocols such as P2, P3, and P4, one could overcome differences in mappability by deeper sequencing (e.g., >2X deeper, **Fig. 3a, b**). Thus, for the WGS of SARS-CoV-2 involving large numbers of samples we believe that using the RNA isolated from frozen samples (−80°C) can be a practical and better choice.

The ARITIC amplicon-based target whole-genome amplification of SARS-CoV-2 is considered as a highly sensitive and low-cost method which could provide high coverage for the viral genome with much less sequencing needed^24^. Several studies have used the ARTIC target whole-genome amplification technique for sequencing SARS-CoV-2^14, 25, 26^. The QIAseq SARS-CoV-2 Primer Panel protocols P1 and P7 were based on the ARTIC V3 primer set, but with a replacement of the 76_RIGHT primer by a substitute primer (i.e., 5′-TCTCTGCCAAATTGTTGGAAAGGCA-3′)^27^. Consistent with the previous reports^28^, our study showed that P1 and P7 preferentially amplified SARS-CoV-2 genome up to 100-fold over human or bacterial genomes in human samples (**Fig. 2d-f**). Compared to the RNA-seq metagenomics-based technologies (i.e., P2, P3 and P4), P1 and P7 achieved more than 100-fold higher coverage for the SARS-CoV-2 genome depending on viral load and sequencing depth (**Fig. 2d-f**). At high viral input, as few as 50K reads were sufficient for P1 and P7 to achieve > 90% viral genome coverage (min10X) (**Fig. 3b**). We found that the P7 worked better than P1 for the samples with low viral copy number or even for samples with partial RNA degradation (**Fig. 3a** **and Suppl. Figs. 3a & 5**). Furthermore, we also noticed that P1 had noticeably more bias and large variations (spikes) in genomic coverage of several regions which were associated with the primer sets 19-21, 43-47, 75-77, and 88-90, respectively. However, these variations were significantly decreased with P7, which showed a much more uniform genome coverage at both low and high viral inputs (**Fig. 4c**, **Suppl. Fig. 4&5**).

Although the primer-panel based target amplicon sequencing has been shown as a cost-effective approach for sequencing the clinical COVID-19 samples to discover the individual genetic diversity^29^, we found there were some limitations for the ARTIC V3 amplicon-based target whole-amplification protocols. First, by design, the current ARTIC V3 amplicons only covered genome regions from positions 30 to 29836, which would make it impossible for the ARTIC V3 amplicon-based protocols to detect a SNV outside of the PCR amplified regions. This scenario actually occurred in our benchmarking study and we found that a consensus variant, g.29868 G>A in sample NP29, was consistently detected by protocols P2, P3, and P4, but was missed by P1 and P7 (**Fig. 5a, c&d**). Second, a single-base mismatch between the primer and template may produce a PCR error such as chimeric PCR amplification^30^, which might lead to a false SNV call. For example, we found that P7, at low viral input, called a unique “false” SNV (g.28321 G>T) with almost 100% allele frequency and >1,000X coverage (**Suppl. Fig. 13)**. However, this putative SNV was not detected in the same clinical sample prepared using either P7 at high viral input (1M) or P1, P2, P3 and P4 (**Fig. 5c**, **Suppl. Fig. 13**) at any input. Third, PCR amplified primer-originated “contaminated” sequences associated with the Qiagen protocols P1 and P7 may lead to an error in SNV calling. Coincidently, we had a consensus SNV (g.6543 C>T) which was within the overlapping binding site to the right adjacent primers 21 and 21alt. Interestingly, this SNV was consistently called by P2, P3, P4, and P7 at the defined threshold (>80% frequency), but in P1 had a significantly lower variant allele frequency (**Fig. 5a&c**) and was not called. Because the PCR primers could mask a SNV that was located in the primer-binding regions, proper primer trimming would be critical for accurately detecting SNVs within the primer binding regions. To understand how this inconsistency occurred, we analyzed the sequencing reads derived from the amplicons 21 and 22 (after adapter trimming) before and after primer trimming on the sequencing data generated from both P1 and P7. We found that, for P1, after adapter trimming, only 7.18% of the reads contained g.6543 C>T SNV; but after primer trimming using either iVar or CLC (Qiagen, https://www.qiagenbioinformatics.com/products/clc-genomics-workbench), the frequency of g.6543 C>T became 43.09% or 6.46%, respectively, whereas many reads containing the primer sequences (g.6543 C) still remained and were not trimmed properly (**Extended Data Fig. 1a**). For P7, after adapter trimming, 63.03% of the reads contained the consensus g.6543 C>T SNV; after primer trimming using either iVar or CLC, the frequency of g.6543 C>T became 81.7% or 62.42%, respectively (**Extended Data Fig. 1b**). Per Qiagen protocols, during the library constructions for P1 and P7, the PCR amplified products were subject to an enzymatic random fragmentation which could generate primer-originated “contaminated” sequences, i.e., the reads containing partial primer sequences or reverse complementary complete/partial primer sequences that could not be removed by iVar or CLC (**Suppl. Fig. 14**). However, when applying Cutadapt^31^, a trimming algorithm that removed all the partial or complete primer sequences by trimming only the end of the reads, i.e., “end-primer sequence trimming”, the frequency of the SNV calling for g.6543 C>T became 91.14% (P1) or 95.79% (P7), respectively (**Extended Data Fig. 1**).

In contrast to the ARTIC V3 amplicon-based target genome amplification, the RNA-seq based metagenomics sequencing protocols such as P2, P3 and P4 used an unbiased approach to cover the whole-genome. The metagenomics approach has been used for sequencing SARS-CoV-2 in several recent studies^10,11,13,32^. Obviously, a unique advantage of the metagenomic approach is its whole-genome coverage including all bases for the SARS-CoV-2 genome given an adequate sequencing depth. We found that when the samples contained a higher viral load (e.g., ~250K or 1M copies), P2, P3, P4 achieved almost complete SARS-CoV-2 genome coverage (min10X) with only ~1M reads per sample (**Fig. 3b**). When the samples NP08, NP29, and NP30 contained a lower viral input (< 1K copies), we observed only P2 achieved sufficient whole-genome coverage (min10X, **Fig. 3a**) leading to 100% sensitivity for detecting the consensus SNVs at a depth of ~5M reads (**Suppl. Fig. 9**). The protocol P4 is based on single primer isothermal amplification technology (SPIA, NuGEN) coupled with the high-throughput sequencing, which can also generate a full-length SARS-CoV-2 genome. The SPIA has been shown to generate the full-length genomes for HIV, West Nile virus, and bovine coronavirus etc^33,34^. However, for samples with low viral input (< 1k copies), we observed that only 0.003% of the SPIA reads could map to the viral genome, suggesting that P4 might not be ideal if a low copy number of SARS-CoV-2 within sample is expected.

Detecting individual SARS-CoV-2 genome variation is critical in tracking the viral spread, evolution, as well as for understanding the potential drug resistance. Thus, we benchmarked and ranked the sensitivity, reproducibility, and precision of the SNV calling of SARS-Co-V-2 across protocols (**Extended Data Fig. 2** **and Suppl. Fig. 15**). We found that the metagenomics protocol P2 was ranked consistently best in the sensitivity of SNV detection, followed by P7, P3 and P1 at either low or high viral input (**Extended Data Fig. 2a** **and Suppl. Fig. 15a**). The rankings for reproducibility of SNV calling were very similar to rankings for sensitivity of SNV calling, although differences between top protocols were smaller and P4 at high viral input moved up in rank. (**Extended Data Fig. 2b** **and Suppl. Fig. 15b**). In contrast, there was a striking difference in the ranking order between the low viral input and high viral input regarding precision, i.e., P7 was ranked best followed by P3 and P1 at low viral input; whereas at high viral input, P7 and P2 performed the best, followed by P3, P1, and P4 (**Extended Data Fig. 2c** **and Suppl. Fig. 15c**). However, at low viral input, all protocols including P7 performed poorly for precision, thus the ranking may be somewhat random. At high input, the order is probably more meaningful. Therefore, we should not be surprised if the precision ranking for some protocols changes dramatically between low and high inputs. Overall, we observed that the viral input was a key factor impacting the SNV calling sensitivity, reproducibility, and precision for SARS-CoV-2, e.g., a low viral input adversely affected the SNV detection (**Figs. 5a-b**, **6a-b**, **Extended Data Fig. 2a-c**, **Suppl. Figs. 8, 9 & 15**). As expected, limited copy number of the viral RNA requires extra PCR amplification, which inevitably introduces more noise and bias^35^ as well as potential errors^30^. Other studies have reported similar effects of viral input copy number on the sequencing and mutation detection quality in line with our observations^16,36–38^.

We also ranked the protocol performance based on mappability, minimal genome coverage and uniformity of genome coverage of the SARS-Co-V-2 (**Extended Data Fig. 2d-f** **and Suppl Fig. 15d-f**). Obviously, P7 and P1 performed much better than metagenomics protocols P2, P3, and P4 in the mappability at both low and high viral input (**Extended Data Fig. 2d** **and Suppl. Fig. 15d**). In addition, P7 was ranked best uniformity of genome coverage at both low and high viral input, whereas P3, P4 and P2 also performed generally well (**Extended Data Fig. 2f** **and Suppl. Fig. 15f).** However, for minimal genome coverage (% of genome with min10X), metagenomics protocols P2, P3 and P4 consistently outperformed the primer-panel based protocols P7 and P1 at both low and high viral inputs (**Extended Data Fig. 2e** **and Suppl. Fig. 15e)**.

In conclusion, our study shows that metagenomic approaches are more sensitive, reproducible and accurate at moderate to higher read depth (e.g., 5M reads) for the SARS-Co-V-2 SNV calling. Although amplicon approaches produce high coverage at lower read depths, they may yield less accurate detection (more false positives and false negatives), leading to reduced sensitivity compared to other methods for the reasons stated previously. Therefore, for protocols P2-P4, we recommend at least 1M viral copies for input and 5M PE reads so that reasonable levels of sensitivity, reproducibility, and precision are achieved. If lower read depths are preferred, then with P7, one can achieve satisfactory high levels of sensitivity, reproducibility, and precision in SNV calling with fewer reads (0.5M-1M PE reads), especially with a reasonable threshold for allele frequency and if the variants are within the amplicon design. In summary, we benchmarked SARS-CoV-2 whole-genome sequencing using seven NGS protocols and evaluated the differences in mappability, viral genome coverage, and variations in SNV calling sensitivity, reproducibility and concordance across input amounts and between protocols. The result of our study will provide a thorough reference and resource on selecting appropriate whole-genome sequencing technologies for clinical SARS-CoV-2 samples, providing knowledge to mitigate the impact of COVID19 on our society.

## Methods

### Study design

**Figure 1** illustrate our overall study design. Briefly, eight COVID-19 positive nasopharyngeal swab RNA samples, either freshly isolated or from frozen condition, were used to generate SARS-CoV-2 WGS libraries using seven protocols (**Fig. 1**, **Suppl. Table 3**). Two different SARS-Co-V-2 inputs, low (1000 copies) vs. high (250,000 or 1 million copies) were used. Each pair-wise low vs. high input were obtained the same clinical sample except P4 for which the low viral input WGS libraries were obtained from different samples due to the limitation of minimal total RNA amount required (**Suppl. Table 3**). For fresh samples, three different viral inputs, i.e., 1000 vs. either 250,000 or 1 million SARS-Co-V-2 viral copies, were used from each same sample, whereas for frozen samples, two different viral inputs, i.e., 1000 vs. 1 million SARS-Co-V-2 viral copies, each from the same sample were used. The seven protocols included: the QIAseq SARS-CoV-2 Primer Panel V1 ARTIC V3 primer set based target genome amplification protocol (P1); the QIAseq FX Single-cell RNA-seq library kit coupled with QIAseq FastSelect −rRNA HMR kit protocol (P2); the QIAseq FX Single-cell RNA-seq library kit coupled with QIAseq FastSelect -rRNA HMR kit and QIAseq FastSelect −5S/16S/23S kit protocol (P3); the Tecan Trio RNA-seq kit coupled with human rRNA depletion protocol (P4); protocols 5 and 6 used an in-house cDNA synthesis recipe with a mix of random primers, oligo(dT), and four pairs of SARS-CoV-2 specific primers, coupled with either the Illumina DNA library preparation kit-DNA Nano (P5) or the Nextera XT kit (P6); the QIAseq SARS-CoV-2 Primer Panel V2 ARTIC V3 primer set based target genome amplification protocol with proprietary buffer chemistry modification (P7). The SARS-Co-V-2 WGS libraries were sequenced, pair-end, 300×2 or 150×2 bp, on two different Illumina platforms (MiSeqDx vs. NextSeq 550, **Suppl. Table 4**). We benchmarked the performances of protocols on mappability, viral genome coverage (%) and uniformity, and sensitivity, reproducibility, and precision of SNV calling.

### Ethics statement

The study was approved by the Institutional Review Board (IRB number 5200127) and the Institutional Biosafety Committee (IBC) of the Loma Linda University (LLU). All the clinical specimens were collected at LLU Medical Center.

### Clinical COVID-19 specimens and RNA isolation

The nasopharyngeal (NP) specimens were collected from SARS-CoV-2 positive individuals. A total of eight nasopharyngeal swab specimens were included in the evaluation. After collection, the NP swab was immediately immersed in 700 μl diluted AVL buffer (140 μl PBS + 560 μl AVL buffer) in an Eppendorf tube and incubated in room temperature for 10 minutes. For fresh samples, the RNA was extracted within 1 to 2 hours of sample collection. For frozen samples, the tubes were placed in −80 °C for 5-6 days before RNA isolation.

RNA was isolated from NP swabs with QIAamp viral RNA mini kit (Qigen, Germany) according to manufacturer’s instructions, and 5.6 μg carrier RNA in AVE buffer and 560 μl absolute ethanol were added to each sample and mixed well. The entire volume of lysate was passed through QIAamp Mini column and centrifuged at 6,000 g for 1 min, allowing RNA to bind to the column. Following the wash by AW1 and AW2 buffer, the RNA was eluted in 50 μl AVE buffer and stored in −80 °C for down-stream procedures.

### DNase digestion on selected RNA samples

15 μl purified RNA was mixed with 72.5 μl of nuclease-free H2O, 10 μl of Qiagen Buffer RDD, and 2.5 μl of RNase-free DNase I stock solution (Qiagen). The sample was incubated for 10 minutes at room temperature. Then, the RNA was purified using the RNeasy MinElute Cleanup Kit (QIAgen), following manufacture’s protocol.

### SARS-CoV-2 viral load determination by qRT-PCR

To confirm the presence of SARS-CoV-2 RNA and the viral copy number from clinical specimens, real-time RT-PCR was performed using SYBR green qRT-PCR method (**Suppl. Table 2**). The 2019-nCoV primer and probe sets consisted of SARS-CoV-2-specific Orf1ab, spike (S) gene, nucleocapsid (N) gene, and human RNaseP. Synthetic positive control (Applied Biosystems) containing the target sequences for each of the assays included in the 2019-nCoV Assay Kit, was included in each assay. Prior to quantitating SARS-CoV-2 viral load in clinical specimens, amplification efficiencies and limit of detection were assessed using six dilutions of 2019-nCoV positive control. All four RT-PCR reactions were performed for each sample in duplicate using the following cycling conditions on an Applied Biosystems QuantStudio 7 Instrument (Applied Biosystems): reverse transcription at 50 °C for 5 min, initial activation at 95 °C for 20 sec, followed by 40 cycles of 95 °C for 3 sec and 60 °C for 30 sec, during which the quantitation of products (FAM) occurred. The efficiency of all four rRT-PCR reactions were higher than 99% (r^2^≥99.97%) and all three coronavirus reactions (Orf1ab, S, N) were required to be positive (Ct <38), regardless of the quantity of RNaseP detected. A result was deemed negative if all three reactions failed to detect the SARS-Co-V-2 target genes (Ct ≥ 38) but RNaseP was detected (Ct< 35). A result was deemed inconclusive if one or two of the three failed to amplify (Ct ≥ 38). A result was deemed invalid if all three coronavirus reactions were negative and if there was evidence of insufficient input (e.g., RNaseP Ct ≥ 35). The SARS-CoV-2 genome copy number in total RNA was quantitated by comparing the average coronavirus Ct across Orf1ab, S, and N to that of the positive control.

### SARS-CoV-2 WGS library construction using QIAseq SARS-CoV-2 Primer Panel V1 (ARTIC amplicon) library protocol (Protocol 1)

The SARS-Co-V-2 viral target genome amplicon libraries were constructed using QIAseq SARS-CoV-2 Primer Panel V1 (Qiagen, Germany) coupled with QIAseq FX DNA library kit (Qiagen) following the manufacturer’s protocols.

Briefly, 5 μl of total RNA of different viral input (1 million, 250,000 or 1,000 viral copies, respectively) was reverse transcribed to synthesize cDNA using random hexamers. 5 μl of cDNA was evenly split into two PCR pools (2.5 μl for each pool) and amplified into 400 bp amplicons using two sets of primers which cover 99% of the entire SARS-CoV-2 genome. The Qiagen primer panel was designed based on ARTIC V3 primers, with the exception that the right primer for amplicon 76 (nCoV-2019_76_RIGHT(-)) was replaced with a modified sequence, 5′-TCTCTGCCAAATTGTTGGAAAGGCA-3’ (Itokawa group). The PCR was performed per manufacturer instruction with 25-cycle amplification for 1 million and 250K viral copy samples, and 32-cycle amplification for 1,000 viral copy samples. After amplification, the contents of 2 PCR pools were combined into one single tube for each sample followed with an AMPure bead clean-up per manufacturer’s instruction. The purified amplicons were quantified using Qubit 3.0 (Life Technology) and normalized for DNA library construction.

Two inputs (40 ng and 1.8 ng) of purified amplicons were used for DNA library construction. Enzymatic fragmentation and end-repair were performed to generate 250bp DNA fragments with an adenine on the 3’ end. Two different fragmentation time were applied depending on different DNA inputs, i.e., high input (1 million and 250K viral copy samples), 18 minutes; low input (1,000 viral copy samples), 12 minutes. The Illumina adaptors were ligated to the DNA fragments followed by AMPure bead clean up. The AMPure bead cleaned up DNA products were further amplified, i.e., 8-cycles for the 40 ng input of amplicons or 20 cycles for the 1.8 ng input of amplicons. The final libraries were quantified by Qubit 3.0 (Life Technology) and quality analyzed on a TapeStation 2200 (Agilent).

### SARS-CoV-2 WGS library construction using QIAseq FX single-cell RNA-seq library protocols (Protocols 2 and 3)

Two sets of libraries were constructed using the QIAseq FX single cell RNA library kit (Qiagen, Germany), with one set coupled with the QIAseq FastSelect rRNA HMR kit only (Protocol 2), and the other set coupled with both the QIAseq FastSelect rRNA HMR kit and the QIAseq FastSelect bacterial 5S/16S/23S kit (Protocol 3) following the manufacturer’s instructions. For fresh samples, three different viral inputs, i.e., 1000 vs. either 250,000 or 1 million SARS-Co-V-2 viral copies, were used from each sample, whereas for frozen samples, two different viral inputs, i.e., 1000 vs. 1 million SARS-Co-V-2 viral copies, were used.

Briefly, to deplete human ribosomal RNA, 1 μl of diluted (0.08x) FastSelect rRNA HMR (Qiagen, Germany) was added into 6 μl COVID-19 specimen RNA along with 3 μl NA denaturation buffer, followed by heated at 95°C for 3 min and then stepwise cooled to 25°C for 14 min. Afterwards, reverse transcription was performed using both random primer and oligo dT primer, and the remaining library preparation steps were performed following the protocol of QIAseq FX single cell RNA library kit (QIAGEN, Germany).

To deplete both human and bacterial ribosomal RNA, 1 μl of diluted (0.08x) FastSelect rRNA HMR (Qiagen, Germany) and 1 μl of diluted (0.08x) QIAseq FastSelect 5S/16S/23S (Qiagen) were added into 6 μl COVID-19 specimen RNA along with 3 μl NA denaturation buffer, followed by heated at 95°C for 3 min and then stepwise cooled to 25°C for 14 min. QIAseq Bead cleanup was carried out per the manufacturer’s instructions. Afterwards, reverse transcription was conducted using both random primer and oligo dT primer, and the remaining library preparation steps were performed by following the protocol of QIAseq FX single cell RNA library kit (Qiagen).

After REPLI-g amplification, 200 - 1000 ng of input cDNAs were used for enzymatic fragmentation by incubating at 32°C for 15 min, followed by adaptor ligation and AMPureXP bead cleanup. Final libraries were eluted from the beads without amplification. All the libraries were quantified with Qubit 3.0 (Life Technologies) and quality analyzed on a TapeStation 2200 (Agilent).

### SARS-CoV-2 WGS library construction using Tecan Trio RNA-Seq library protocol (Protocol 4)

Eight RNA samples isolated from fresh and frozen specimens were used for Tecan Trio RNA-seq library construction (NuGEN/Tecan), following the NuGEN protocol with integrated DNase treatment. For fresh sample, total RNA amounts containing 1 million or 250,000 SARS-Co-V-2 viral copies for NP08 and NP17, and 1000 SARS-Co-V-2 viral copies for NP12 and NP16 were used as input. For frozen samples, total RNA amounts containing 1 million or 250,000 SARS-Co-V-2 viral copies for NP29 and NP30, and 1000 SARS-Co-V-2 viral copies for NP26 and NP27 were used as initial input, respectively. All procedures were carried out using conditions specified in the Trio RNA-seq protocol.

The 10 μl of total RNA was treated with DNase, followed by cDNA synthesis using random hexamers. After purification by AMPure XP beads (Beckman Coulter), cDNAs were amplified on beads by single primer isothermal amplification (SPIA). Next, enzymatic fragmentation and end repair were performed to the cDNAs to generate blunt ends. The Illumina adaptors were ligated to cDNA fragments, followed by first round of library amplification. AnyDeplete probe mix was used to deplete the human ribosomal transcripts. The remained DNA libraries were amplified a second time for 6 cycles. Additional 9 cycles of amplification were carried out for the libraries with a yield lower than 10 ng.

After the second round of library amplification, double size selection by AMPure beads was performed to obtain library molecules with size ranged between 200 bp and 700 bp. Then, 22.5 μl of AMPure beads was added to the 50 μl library products. After incubation and magnetic separation, supernatant was collected and another 22.5 μl of AMPure beads was added. Following magnetic separation, the supernatant was removed and the beads were washed with 70% alcohol. The final libraries were eluted in water. The libraries were quantified by Qubit 3.0 (Life Technology) and quality analyzed on a TapeStation 2200 (Agilent).

### SARS-CoV-2 WGS library construction using metagenomics approach combining an in-house cDNA synthesis recipe, Qiagen MDA and Illumina DNA library protocol (Protocols 5 and 6)

The amount of RNA input was normalized based on the viral load determined by SYBR green qRT-PCR method. Specifically, total RNA amounts containing 1 million, 250,000 or 1000 SARS-CoV-2 viral copies were used to start the initial cDNA synthesis by SuperScript III reverse transcriptase (Invitrogen), using a mix of random primers, oligo(dT)18, and four pairs of SARS-CoV-2 specific primers that cover the 5’ and 3’ ends. The sequences of the specific primers were: F1, 5′-ATTAAAGGTTTATACCTTCCC-3′; R1, 5′-TTTTTTTTTTTTGTCATTCTCC-3′; F2, 5′-TTCTTATTTCACAGAGCA-3′; R2, 5′-AACATAACCATCCACTGAATATG-3′; F3, 5′-AAATGGGGTAAGGCTAGAC-3′; R3, 5′-AGTCTACTTGACCATCAAC-3′; F4, 5′-AGCACACTTTCCTCGTGAAGG-3′; R4, 5′-CTTGAACTTCCTCTTGTCTG-3′. Reverse transcription (RT) primer annealing was conducted at 65°C for 5 minutes, then, incubated on ice for 1 minute. RT was carried out at 25 °C for 10 min, then 55 °C for 30 min in 10 μL volume. All RT products were used for cDNA amplification using QIAseq multiple displacement amplification (MDA) technology. After amplification, the cDNA was purified using equal volume of Agencout AMPure XP beads (Beckman Coulter). 100 ng or 1 ng purified cDNAs were used for library construction using either TruSeq DNA Nano library preparation kit (Illumina), i.e., P5, or Nextera XT DNA library preparation kit (Illumina), i.e., P6, respectively. All procedures were carried out following the protocols recommended by the manufacturers. The libraries were purified with Agencourt AMPure beads, quantitated by Qubit dsDNA HS assay (Life Technologies), and the quality was analyzed on a TapeStation 2200 with D1000 Screen tape (Agilent).

### SARS-CoV-2 WGS library construction using QIAseq SARS-CoV-2 Primer Panel V2 (ARTIC amplicon) library protocol (Protocol 7)

The SARS-Co-V-2 viral target genome amplicon libraries were constructed using the QIAseq SARS-CoV-2 Primer Panel V2 (Qiagen, Germany) coupled with QIAseq FX DNA library kit (Qiagen) following the manufacturer’s protocols. All the procedures including primers were identical as described in P1, except for some proprietary modifications on the buffers (Qiagen, personal communications).

### SARS-CoV-2 WGS library sequencing

The libraries were multiplexed with different barcodes and pooled at 4 nM in equimolar amounts. The pooled libraries were quantified by Qubit prior to sequencing. The pooled libraries were clustered on Illumina NextSeq 550 high output flow cell at a final concentration of about 2.1 pM and MiSeqDx flow cell at 8.5 pM. The libraries were sequenced on an Illumina NextSeq 550, pair-end, 150×2 bp. The same libraries were also sequenced on an Illumina MiSeqDx, pair-end, 300×2 bp or 150×2 bp (Illumina, Inc., San Diego, CA).

### Sequence data processing, mapping, and mapping rate generation

Sample QC were reported by fastqc^39^, qualimap^40^, and MultiQC^41^. The raw reads were trimmed with cutadapt (v1.9.1). The trimmed reads were aligned to the Wuhan-Hu-1 reference using bwa mem (v0.7.12)^42^ with default settings. For the sequencing data generated using the ARTIC V3 based primer-panel protocols (P1 and P7), an extra primer trimming was performed using iVar (v1.2.2). The aligned reads were further de-duplicated by samtools rmdup (v1.9)^43^ to get the bam files. A Kraken2^44^ database was built based on the complete genomes in the NCBI RefSeq database for archaea, bacteria, protozoa, fungi, human and viruses (SARS-COV-2 genome included). To summarize the read mapping percentages to multiple taxa, the trimmed reads were classified into human, SARS-CoV-2, bacterial, and remaining reads (e.g., unclassified, archaeal, viral, fungi, protozoa) by using the Kraken2 database. The sequencing read mappability (mapping percentage) to the SARS-Co-V-2 genome was computed for each of the 7 protocols at different sequencing depths.

### Statistical analysis of factors impacting mappability

A generalized linear mixed model (glmm) was created using lmer package^45^ in R to determine the significant factors explaining variations in mapping rates. As mapping rates tended to be at extremes for different protocols (i.e., mapping rates were usually near 0 or 100%), we first transformed the mapping rate using the probit transform. The zero value in mapping rate was replaced with 1.00E-09 for the probit transform. We then created a glmm using the transformed mapping rate as the dependent variable and employed fixed effects of protocol, viral copy input amount (low or high), and sample storage condition (fresh vs. frozen) and a random effect of the viral RNA input concentration. No interactions terms were significant and results from the simplest model with these factors were reported. P values were generated using the Satterthwaite approximation for degrees of freedom carried out by the lmerTest package in R. To identify which groups were statistically different from one another, pairwise comparisons were carried out for all the fixed effects using difflsmeans function of lmerTest package^46^ in R. The degree of freedom was adjusted by Satterthwaite method.

To evaluate sequence mapping to SARS-CoV-2 viral, human, and bacterial genomes, data were presented as the mean ± one standard deviation. To evaluate sequence read depth on minimal viral genome coverage, data were presented as the mean ± one standard error.

### Genome coverage and coverage uniformity calculation

The genome coverage was defined as the breath of coverage, which was measured as percentage of the SARS-Co-V-2 reference genome for which the genomic positions (bases) were sequenced with minimal 10× coverage. Coverage uniformity on the SARS-CoV-2 genome was examined by comparing a quantitative metric, i.e., coefficient of variation (CV) across seven protocols. CV was computed using the standard deviation and mean of the coverage at each reference genome position.

### SARS-Co-V-2 SNV variant calling and generation of consensus SNV variants

Variants were called on the bam files by VarScan 2 (v2.4.4) and BCFTools (v1.9). To accurately identify SNVs, we used samtools mpileup (parameters: -A -d 20000 -Q 0) and varscan2 (v2.4.4) (parameters: --p-value 0.99 –variants). Then, we filtered the low-confidence SNVs with snippy vcf_filter (parameters: --minqual 100 --mincov 10 --minfrac 0.8).

Samtools (v1.9) mpileup and BCFTools (v1.9) were used to generate the genome variants fastq file from the bam file, and the fastq file was converted into genome fasta file using Linux cat command. The variant fasta files generated from the same clinical sample but different protocols were piled up using Jalview (v2.11.1.0)^47^ alignment tool, from which one consensus fasta file was compiled for each clinical sample.

### Calculations of sensitivity, reproducibility, and precision of SNV calling

#### Sensitivity

Variant calls from iVar were filtered for variant allele frequency or VAF (only putative variants with VAF > 80% were considered as called variants). Sensitivity used the consensus SNVs (discussed earlier) as the set of positives. Sensitivity was defined as TP/(TP + FN) (TP, true positive; FN, false positive). Protocols were ranked by average sensitivity using 5M reads for all protocols except P7, where 1M reads were used (P7 was not sequenced at 5M reads).

#### Reproducibility

Variant calls from iVar were filtered for variant allele frequency. For **Figure 6**, multiple VAF (0.1, 0.5, 0.8, 0.9) thresholds were considered for illustration (as well as different input levels and read depths) to establish variant calls by sample and by protocol. Given a set of variant calls, reproducibility between hypothetical protocol A and one of the other protocols was defined as the percent of variants identified by protocol A at VAF > 0.8 that was also identified by the other protocol at any allele frequency for each sample. At low input, each protocol had three other protocols with which to compare (P4, P5, and P6 did not have calls at low input) with three samples, yielding nine data points for each protocol. At high input each protocol had four other protocols with which to compare (P5 and P6 did not have calls at low input) with three samples, yielding 12 data points for each protocol. Protocols were ranked by average reproducibility using 5M reads for all protocols except P7, where 1M reads were used (P7 was not sequenced at 5M reads).

We defined reproducibility relative to an allele frequency threshold: a variant was reproducible between protocols A_i_ and A_j_ if the variant from protocol A_i_’s library had an allele frequency equal to or greater than a threshold and that variant was also identified by protocol A_j_’s library at any allele frequency. For the summary measures of reproducibility between protocols, we averaged reproducibility values across A_i_ using the Jaccard index for reproducibility scoring.

Avg Reproducibility 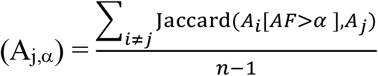 where n is the number of protocols and A_i_ and A_j_ are two sets of called SNVs.

#### Precision

Variant calls from iVar were filtered for variant allele frequency (putative variants with VAF > 20% were considered as called variants, a lower threshold than that used for sensitivity, to better measure potential false calls). Precision used the consensus SNVs (discussed earlier) as the set of positives. Calls of variants with VAF > 20% that were not a consensus SNV were termed false positives (FP). Precision, also known as positive predictive value (PPV), was defined as TP/(TP + FP). Protocols were ranked by average precision using 5M reads for all protocols except P7, where 1M reads were used (P7 was not sequenced at 5M reads).

### Phylogenetic analysis of SARS-Co-V-2 variants

Phylogenetic analysis was performed using Nextstrain pipeline (https://github.com/nextstrain/ncov)^17^ on a local Linux machine. Briefly, five SARS-CoV-2 consensus sequences from LLU NP08, NP12, NP17, NP29, and NP30 samples were combined with the public SARS-CoV-2 genome data (N = 417) from 15 countries downloaded from GISAID (http://gisaid.orgon) on August 15, 2020. The Nextstrain pipeline was run using the combined 417 public and 5 LLU SARS-CoV-2 WGS sequencing data. The output JASON file from Nextstrain pipeline was viewed using auspice (https://auspice.us)^17^ (**Suppl. Fig. 11 and Fig. 12**).

### Bioinformatics analysis for revealing a SNV masked by amplicon primers

In amplicon sequencing, a potential SNV allele could be located within an amplicon primer per se (masked SNV allele) and the amplified viral allele (potential SNV) from an adjacent second amplicon. We had a SNV, i.e., g.6543 C>T (in NP08), which was covered by both n2019_21_R and n2019_21_R_alt primers. The full primer sequence could be removed from the end of a read by standard trimming or by iVar if it was located in anywhere in a read. In both scenarios, the masked SNV allele reads would be removed and the potential SNV could be detected. However, P1 and P7 employed an enzymatic fragmentation step, which resulted in partial primer sequences at the end of some reads that could not be removed by either iVar or CLC Bio package. In addition, iVar may also remove the primer sequences located within the middle of a read, i.e., a “true” SNV derived from the reads amplified by a second adjacent amplicon. Under both circumstances, the potential SNV calling would be compromised. To reveal the masked SNV calling, we compared the g.6543 allele frequencies using different primer trimming methods on fastq file of NP08 at 1M viral input with 5M read depth. Briefly, the same fastq file was trimmed by quality trimming only, CLC Bio (default settings), iVar (default settings), and Cutadapt (ends trimming only), respectively. After trimming, the occurrences of full and partial sequences for primer n2019_21_R and n2019_21_R_alt, as well as their reverse complementary sequences that covered g.6543 were counted and the frequency of allele T was used to compared the efficiencies of trimming methods (**Extended Data Fig. 1**).

### Methods used for protocol ranking

The ranking performances of SNV detection and viral genome mapping across seven protocols were evaluated individually for each of six categories or metrics, using either Z-score statistic based on harmonic mean (**Extended Data Fig. 2**) or displayed by individual sample/data point values (**Suppl. Fig. 15**), both derived from SNV calling and viral genome mapping data at low and high viral input. For the Z-score based rankings, as each protocol contained multiple data points linked to different samples and read depths, a mean was taken as the initial ranking value for the given protocol and was used for Z-score transformation. The Z-score was calculated based on the average of all sample/data points per metric per protocol for either low or high input. To reduce the variation associated with viral input, only data points generated from 1K and 1M viral inputs were used for ranking. Sample NP08, NP29, and NP30 were used for SNV calling ranking evaluations on sensitivity, reproducibility, and precision. Other samples were also included in the mapping-based ranking evaluations on mappability, genome coverage, and uniformity of coverage. SNV detection sensitivity was measured by the percentage of consensus SNVs detected by a protocol at 1M and 5M PE reads. Reproducibility was measured between protocols at low and high input levels indicating whether the SNV calls using a VAF threshold of 80% were reproduced in a different protocol. There was one value per sample per protocol pair (P4, P5, and P6 were not used for low viral input results; P5 & P6 were not used for high viral input results). All data were based on sample sequencing output from 5M PE reads except for P7 which did not have sequencing data at 5M depth thus 1M PE read depth were used instead. Precision of SNV detection was measured with a VAF detection threshold of 20% or more using consensus SNVs as ground truth. Mappability was measured by the percentage of reads that aligned to the SARS-CoV-2 genome and was estimated overall, but not with respect to a particular read depth. The SARS-CoV-2 genome coverage was evaluated based on proportion of the viral genome which achieved at least 10X coverage from both 1M and 5M PE reads in each sample. For uniformity ranking evaluation, the reciprocal values of coefficient of variation (CV) on genome coverage from all samples for each given protocol were included to compute the mean, which was used to calculate Z-score (**Extended Data Fig. 2**). For the rankings displayed based on individual sample or data point for each of six metrics, all sample/data point values were the same as used for the Z-score based rankings, but no mean was calculated, which thus allowed to display the distribution of all samples and data point values for each of the six metrics at low and high viral input (see figure legends for detail in **Suppl. Fig. 15**).

## Data availability statement

The sequencing data have been uploaded to the NCBI SRA (Sequence Read Archive) under the BioProject accession # PRJNA638938. The data will be available to the public when the paper is published. For reviewers, a token for accessing the data can be obtained from the editor of the journal.

Reviewers’ link: https://dataview.ncbi.nlm.nih.gov/object/PRJNA638938?reviewer=r95nj68tijbk8p325rnsddc96n

## Software and code availability statement

We used many algorithms and code sets for the SARS-CoV-2 genome mapping, genome coverage and SNV calling which have been published previously. All of our code is provided in the GitHub at the following link.

https://github.com/oxwang/COVID19_MS1

## Author contributions

CW conceived and designed the study. TL, ZC, and WC performed experiments including RNA isolation, qRT-PCR, SARS-CoV-2 WGS library construction and sequencing. WC, TL, ZC, and CW drafted the manuscript. ZC, TL, WC, XC, WJ, MH, and ZY performed bioinformatics data analysis. WJ and TL conducted biostatistics tests and analysis. WC, ZC, TL, XC, and WJ prepared the methods for the manuscript. DT, CG, and DH helped the patient nasopharyngeal swab sample collection. DH helped coordinate the project meetings and IRB application. CW, WJ, JL, DH, DT, and CG helped revise the manuscript. All authors reviewed the manuscript. CW and WJ edited the manuscript. CW finalized and submitted the manuscript.

## Acknowledgements

The authors are very grateful to Elder Michael Wan, Minister Tom Tui, Ms. Cherry Ngan of the House of Joy Christian Church; and Pastor Ray Yang of the Harvest Chinese Christian Church; for their generous donations of personal protective equipment for our study. The authors would like to thank the LLU Institutional Review Board Committee, particularly Mrs. Amy Casey and Dr. Travis Losey, for their expedited assistance in reviewing of the IRB application. The authors would also like to thank the following people and organizations for the assistances either in securing some critical reagents needed or providing technical consultation to our study during the COVID-19 pandemic, including Ms. Elizabeth Conzevoy, Dr. Sujash Chatterjee, Dr. Jonathan Shaffer, and Mr. Brian M Dugan from Qiagen; Mr. Andrew Shaver, Mrs. Sinem Taylor, and Dr. Denise Stephens from Tecan Genomics. The genomic work carried out at the LLU Center for Genomics was funded in part by the National Institutes of Health (NIH) grant S10OD019960 (CW), the Ardmore Institute of Health grant 2150141 (CW) and Dr. Charles A. Sims’ gift to the LLU Center for Genomics.

## Figures and figure legends for main figures and extended data figures

**Extended Data Figure 1.**
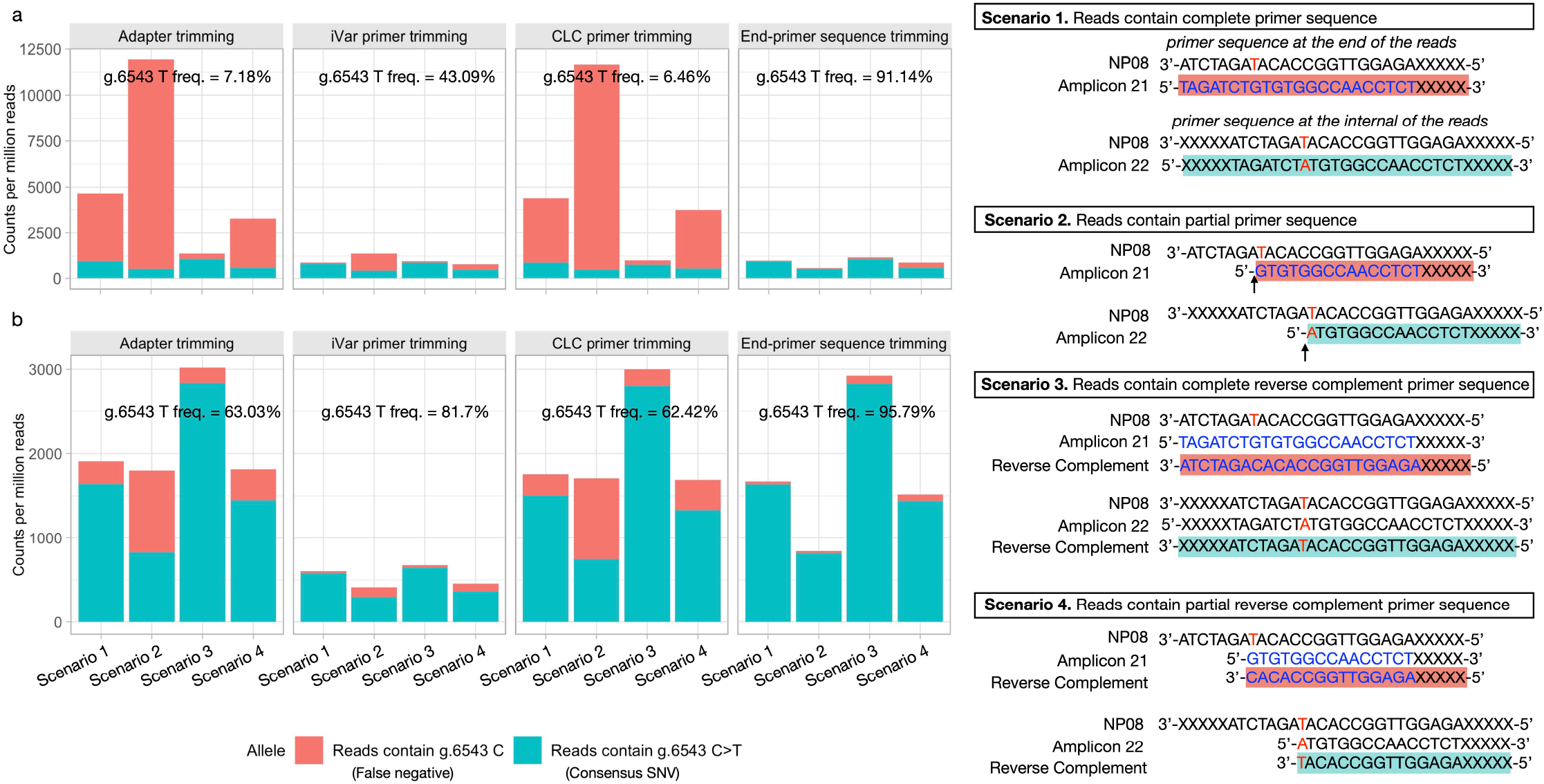
ARTIC V3 primer sequences masked a SNV call of g.6543 C>T. A SNV in NP08 was found in the genome location at g.6543, which was within the primer n2019_21_R and n2019_21_R_alt sequences, and was also in the middle of amplicon 22. After sequencing, reads with g.6543 could be covered by n2019_21_R and n2019_21R_alt sequences under 4 scenarios (**right panel**). Scenario 1: full primer sequences of n2019_21_R and n2019_21R_alt; Scenario 2: partial primer sequences of n2019_21_R and n2019_21R_alt; Scenario 3: full reverse-complementary sequences of n2019_21_R and n2019_21R_alt; Scenario 4: partial reverse-complementary sequences of n2019_21_R and n2019_21R_alt. Fastq files were first trimmed to remove adapters using Cutadapt with default settings, then subject to second round of trimming to remove SARS-CoV-2 ARTIC V3 amplicon primers, using iVar, Qiagen CLC package, and customized Cutadapt trimming. After trimming, the occurrence of primer sequences listed in each of the above scenarios were counted, in searching for evidence that g.6543 C>T SNV call was compromised by primer derived sequence “contamination”. Data was normalized to the number of counts per million reads. (**a**) Library prepared with P1 from NP08 at 1M viral input. (**b**) Library prepared with P7 from NP08 at 1M viral input. Y-axis shows the CPM reads for P1 (**a**) or P7 (**b**); X-axis shows reads containing g.6543 C allele in orange (false negative) and reads containing g.6543 C>T allele in green (consensus true SNV) in four different scenarios (illustrated in the right panel) using four different trimming methods (adapter trimming, iVar primer trimming, CLC primer trimming and non-internal primer trimming). Note that after “non-internal primer trimming”, by applying Cutadapt, a trimming algorithm that removed all the partial or complete primer sequences by trimming only the end of the reads, i.e., “end-primer sequence trimming”, the frequency of the SNV calling for g.6543 C>T became 91.14% (P1) or 95.79% (P7), respectively.

**Extended Data Figure 2.**
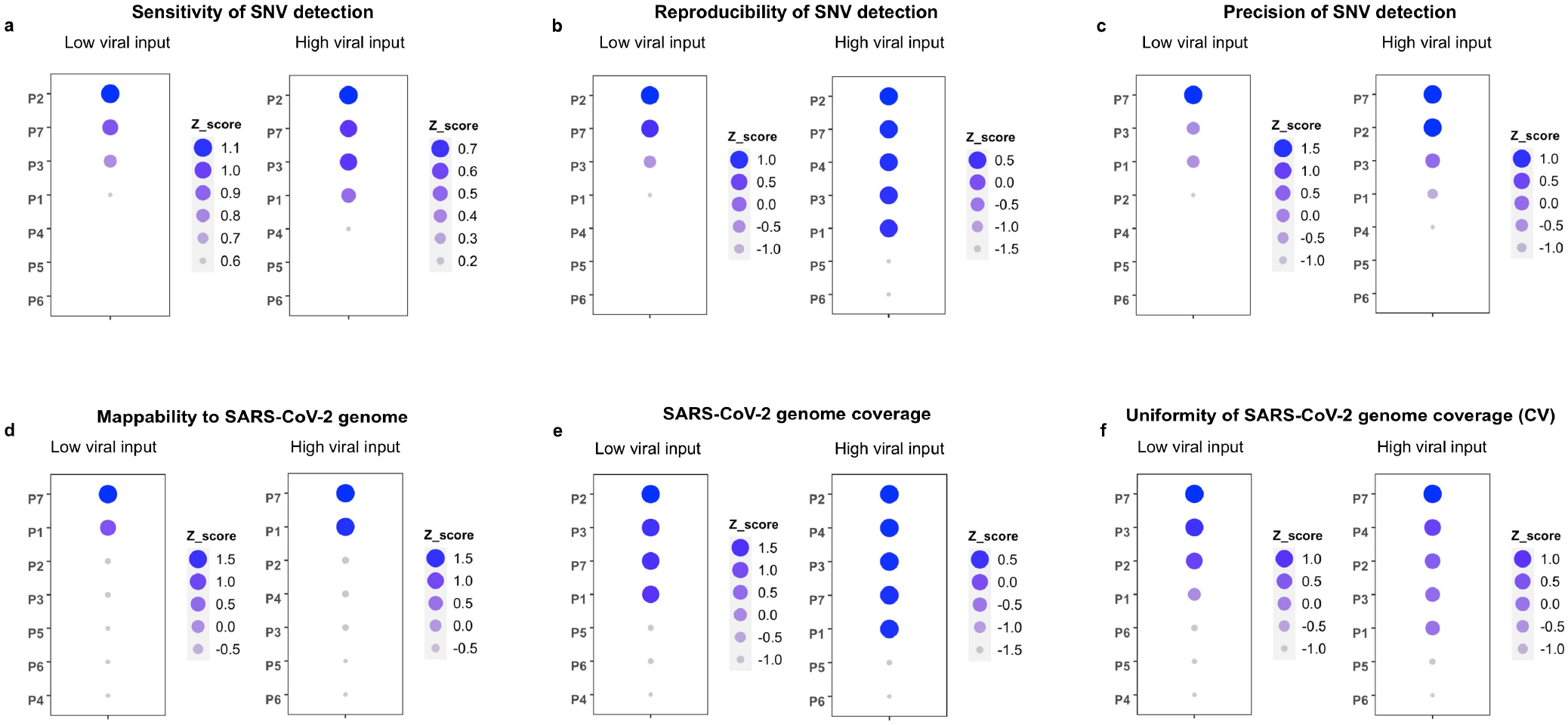
Z-score rankings of SARS-Co-V-2 whole-genome sequencing protocols. Protocols were ranked individually using z-score at each metric. (**a**) Ranking of sensitivity of SNV detection (low and high inputs); sensitivity evaluates the ability of a protocol in detecting potential SNVs based on the consensus SNVs defined (see Results and Methods); (**b**) Ranking of reproducibility of SNV detection (low and high input); reproducibility metric measures the likelihood of SNVs detected in a given protocol that might be detected by another protocol in an independent experiment. Reproducibility and its calculation were defined in the Methods. (**c**) Ranking of precision of SNV detection (low and high inputs); precision metric measures the accuracy of consensus SNV detected by a protocol from all potential SNVs (frequency > 5%); (**d**) Ranking of SARS-CoV-2 genome mappability which measures the mapping efficiency of sequencing data to the viral genome; (**e**) Ranking of SARS-CoV-2 genome coverage which measures the proportion of viral genome that can be covered at specific read depth; (**f**) Ranking of uniformity of genome coverage which evaluates the evenness of coverage across viral genome. The reciprocal value of coefficient of variation (CV) was used for Z-score calculation in order to keep same ranking directionality (large value for better performance) as in other categories. Z-scores are plotted as circles with their size and color shade scaled to the Z-score value from large to small, and dark blue to light blue. Note that larger Z-score values imply better performance,

